# Deep transcriptome sequencing of subgenual anterior cingulate cortex reveals disorder-specific expression changes in major psychiatric disorders

**DOI:** 10.1101/598649

**Authors:** Nirmala Akula, Stefano Marenco, Kory Johnson, Ningping Feng, Joanna Cross, Bryce England, Sevilla Detera-Wadleigh, Qing Xu, Pavan K. Auluck, Kwangmi Ahn, Robin Kramer, Jose Apud, Brent T. Harris, C. Harker Rhodes, Barbara K. Lipska, Francis J. McMahon

**Affiliations:** Human Genetics Branch, and National Institute of Mental Health Intramural Research Program NIH, DHHS, Bethesda, MD USA; Human Brain Collection Core, National Institute of Mental Health Intramural Research Program NIH, DHHS, Bethesda, MD USA; Bioinformatics Section, National Institute of Neurological Disorders and Stroke, NIH, DHHS, Bethesda, MD USA; Georgetown Brain Bank, Histopathology and Tissue Shared Resource, Georgetown University Medical Center, Washington, DC, USA

## Abstract

How do differences in onset, symptoms, and treatment response arise between various mental illnesses despite substantial overlap of genetic risk factors? To address this question, we carried out deep RNA sequencing of human postmortem subgenual anterior cingulate cortex, a key component of limbic circuits linked to mental illness. Samples were obtained from 200 individuals diagnosed with bipolar disorder, schizophrenia, or major depression, and controls. Differential expression analysis in cases versus controls detected modest differences that were similar across disorders, although transcript-level differences were more pronounced. Case-case comparisons revealed greater expression differences between disorders, including many genes and transcripts that were expressed in opposite directions in each diagnostic group, compared to controls. Relative transcript abundances were associated with common genetic variants that accounted for disproportionate fractions of diagnosis-specific heritability. Inherited genetic risk factors shape the brain transcriptome and contribute to diagnostic differences between broad classes of mental illness.

## INTRODUCTION

Genome-wide association studies (GWAS) have revealed that major psychiatric disorders share many common alleles with small additive effects on risk^1,2^. Similarly, gene expression studies in post-mortem brain tissue have found that expression changes in mental illnesses tend to be small and correlated across diagnoses. One large post-mortem study^3^ found small changes in gene expression in schizophrenia (SCZ) versus controls that seemed to reflect the small frequency differences of common alleles associated with SCZ by GWAS^4^. A meta-analysis of microarray expression data drawn from several post-mortem studies found that many differentially-expressed (DE) genes were shared across major psychiatric disorders such as SCZ, autism, and bipolar disorder (BD)^5^. A large follow-up study that used RNA sequencing confirmed substantial overlap in DE genes in SCZ, BD, and autism spectrum disorder, versus controls, although transcript (isoform) level expression was more variable^6^.

If genetic risk factors and gene expression changes in brain are as similar across different mental illnesses as these studies seem to suggest, then how do diagnostic differences in onset, symptoms, and treatment response arise at all? One possibility is that diagnostically distinct genetic risk factors account for some of these differences. For example, genetic correlations as high as the 70% reported between SCZ and BD^1^ are consistent with ~50% of risk alleles differing between disorders. It is also possible that published gene expression studies have overestimated the degree to which gene expression changes in post-mortem brain are similar across disorders. Even the largest sample sizes have been underpowered to detect subtle gene expression differences^3^, and those differences that have been detected may not be representative. Case-control study designs limit opportunities to detect true differences between case groups, and few published post-mortem gene expression studies have focused on direct, case-case comparisons between disorders. Moreover, few if any human post-mortem studies have used methods that reveal most of the transcriptional complexity of the brain, where numerous gene isoforms are generated by alternative splicing and other mechanisms. How much of the apparent resemblance in gene expression reported among clinically distinct mental illnesses can be explained by these methodological limitations?

To address this question, we performed deep sequencing of the brain transcriptome followed by case-control and case-case comparisons of both gene and isoform expression in tissue obtained post-mortem from 200 individuals diagnosed with BD, SCZ, major depressive disorder (MDD), or no known psychiatric illness. Tissue was dissected from subgenual anterior cingulate cortex (sgACC), a key component of limbic circuits that have been implicated in major mood disorders and other mental illnesses^7,8^. Total RNA was sequenced at an average depth of ~270 million reads per sample, generating high-quality expression data for ~21,000 genes and ~85,000 transcripts in 185 samples. We believe these data represent the most complete sampling of the human brain transcriptome published to date.

The results demonstrate that subtle differences in gene expression observed between broad diagnostic classes of mental illness actually reflect more pronounced and diagnosis-specific changes at the transcript level, and that these changes are shaped by inherited genetic risk factors.

## RESULTS

### Transcriptome Profile of the Subgenual anterior cingulate cortex (sgACC)

Total RNA from 200 postmortem brain samples (Controls=61, SCZ=46, BD=39, MDD=54) was captured using the Ribo-zero protocol (see Methods) followed by stranded, paired-end sequencing. The samples, along with demographic information and known covariates are presented in Supplementary Table 1. After quality control, 185 samples were retained for analysis (see Methods). The analysis pipeline is presented in Supplementary Figure 1.

In an effort to detect a more complete set of genes and transcripts than previous studies, RNA sequencing was performed at very high depth, generating ~50 billion reads at an average of ~270M reads/sample. Of these, 167M reads/sample mapped to ~21,000 Ensembl genes and ~85,000 Ensembl transcripts; identifying ~50% more transcripts than those used in previous studies^6,9,10^. The mapping summaries are presented in Supplementary Table 2. Very highly expressed transcripts with >10,000 reads/transcript comprised 1% of the total reads, while transcripts with <100 reads comprised 56% (Supplementary Figure 2). We detected an average of 5 isoforms/gene. The greatest number of isoforms (49 each) were detected for the genes *MUC20-OT1* (a non-coding RNA) and *TCF4*, which has been associated with SCZ, autism, and intellectual disability^11–13^.

### Gene-level Analyses

#### Case-Control comparisons

To explore case-control differences in gene expression, all 21,000 genes were quantile normalized for library size, gene GC-content and gene length. Differential expression analysis was performed for each diagnosis group versus controls using DESeq2^14^, with covariates for RNA integrity number (RIN), reported race, and sample GC content – the only measured variables that were significantly associated, after Bonferroni-correction, with one or more principal components of gene expression (Supplementary Table 3). Sensitivity analyses carried out with surrogate variables for high-dimensional data sets (SVA)^15^ and with additional covariates found similar results, but the chosen 3-covariate model produced the smallest genome-wide inflation in test statistics (Supplementary Figure 3).

As expected, differential expression between cases and controls was modest. Absolute log2 fold change (FC) differences were ≤0.5 in all diagnostic groups. At FDR <5%, a total of 23, 42, and 7 genes were DE in SCZ, BD and MDD, respectively (Figure 1a). Genes that were DE at FDR<5% in more than 1 diagnostic group are shown in Table 1. Many of the same genes were DE in multiple disorders, with a mean overlap of 42% (Figure 2a.). All DE genes with nominal p<0.05 are shown in Supplementary Tables 4, 5 and 6.

**Table1:** Genes Differentially Expressed (FDR <5%) in >1 Diagnostic Group

| Diagnostic group | Ensemble gene ID | Gene name | Direction of change | Gene description | Significance of overlap |
| --- | --- | --- | --- | --- | --- |
| All 3 groups | ENSG00000064300 | *NGFR* | Down | Nerve Growth Factor Receptor | NA |
|  | ENSG00000171551 | *ECEL1* | Down | Damage Induced Neuronal Endopeptidase |  |
|  | ENSG00000179869 | *ABCA13* | Down | ATP Binding Cassette Subfamily A Member 13 |  |
|  | ENSG00000256861 | *AC048338.1* | Down | paralog of VPS33A |  |
| BD and SCZ | ENSG00000172987 | *HPSE2* | Down | Heparanase 2 | $2.22 \times 10^{-14}$ |
|  | ENSG00000175793 | *SFN* | Up | Stratifin, epithelial cell marker protein |  |
| BD and MDD | ENSG00000146066 | *HIGD2A* | Down | HIG1 Hypoxia Inducible Domain Family Member 2A | $4.44 \times 10^{-16}$ |

**Figure 1.**
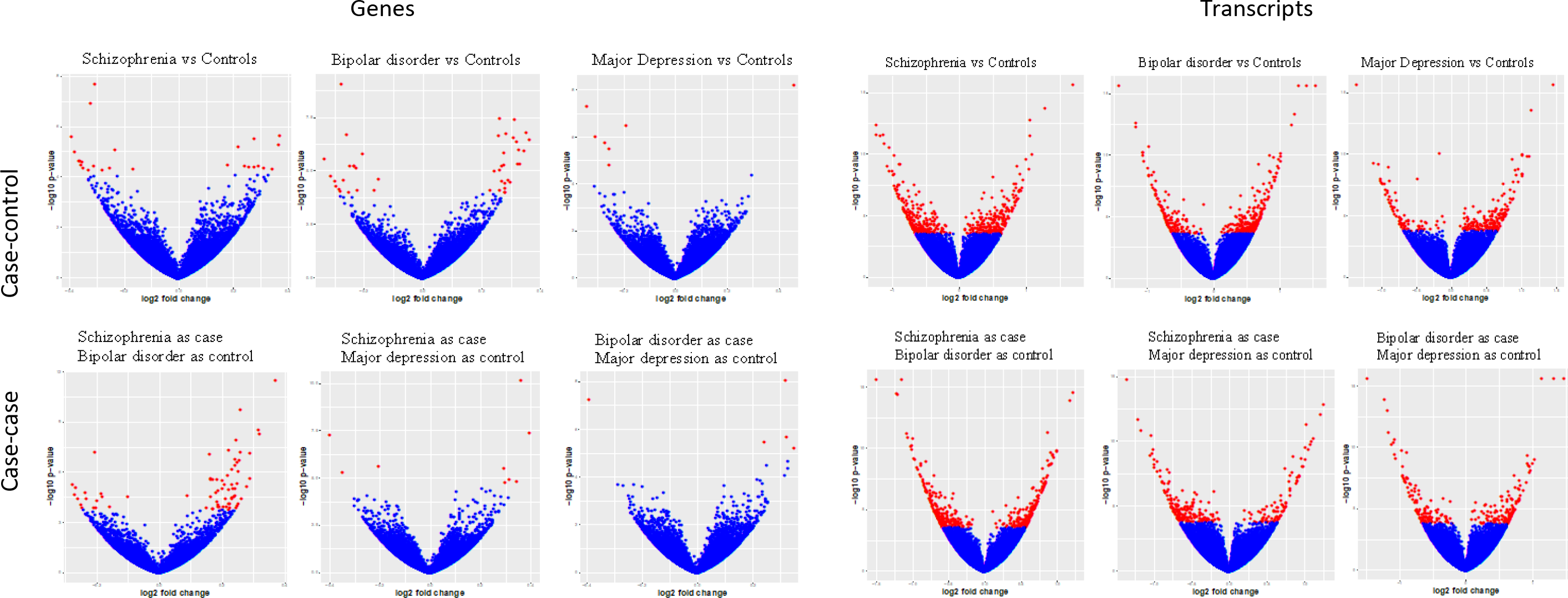
Volcano Plots of Differentially Expressed Genes and Transcripts. Log_2_ fold-change values are plotted against the X-axis and -log_10_ p-values for differential expression are shown on the Y-axis. Genes with FDR<5% are highlighted in red. Gene-level results are shown for a. case-control and b. case-case comparisons; corresponding transcript-level results are shown in panels c. and d., respectively. Note that for case-case comparisons the sign of the log_2_ fold-change is arbitrary.

**Figure 2.**
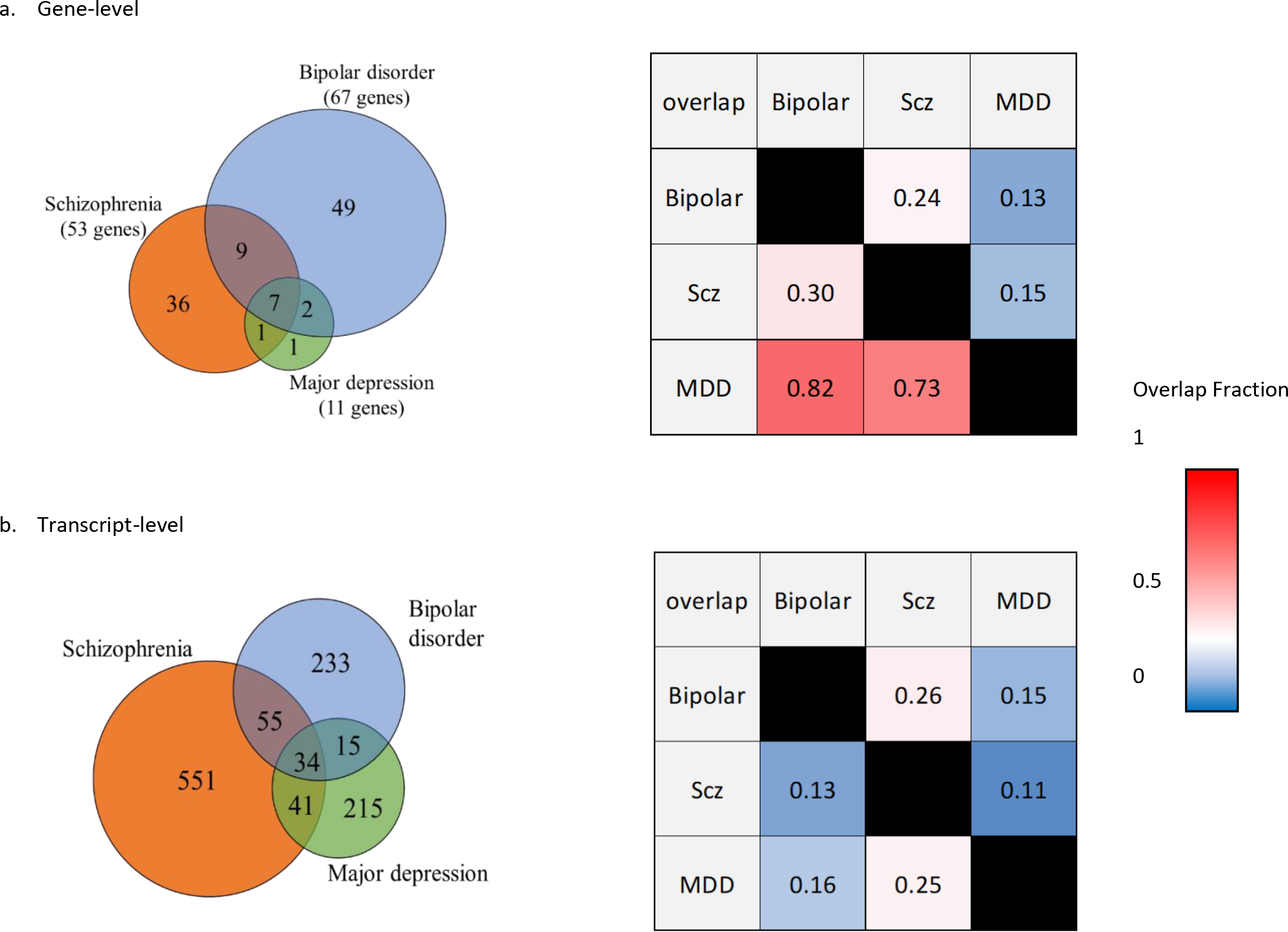
Overlap of differential expression (FDR<5%) across disorders. Gene-level analyses are shown in panel a., with Venn Diagrams of gene name overlaps on the left. Overlaps between diagnostic groups (as a fraction of all differentially-expressed genes) are shown on the right. Corresponding results for the transcript-level analyses are shown in panel b.

Among genes DE within any one diagnostic group, there was a strong positive correlation in FC values across diagnoses (linear regression R^2^ 0.61 to 0.72) (Figure 3a). When we compared the DE genes detected at FDR<5% in sgACC to those identified in a published study of DLPFC^6^, we found significant overlap for SCZ (hypergeometric p-value=8.0×10^−6^) and BD (hypergeometric p-value=6.8×10^−5^) (Supplementary Table 4 & 5).

**Figure 3.**
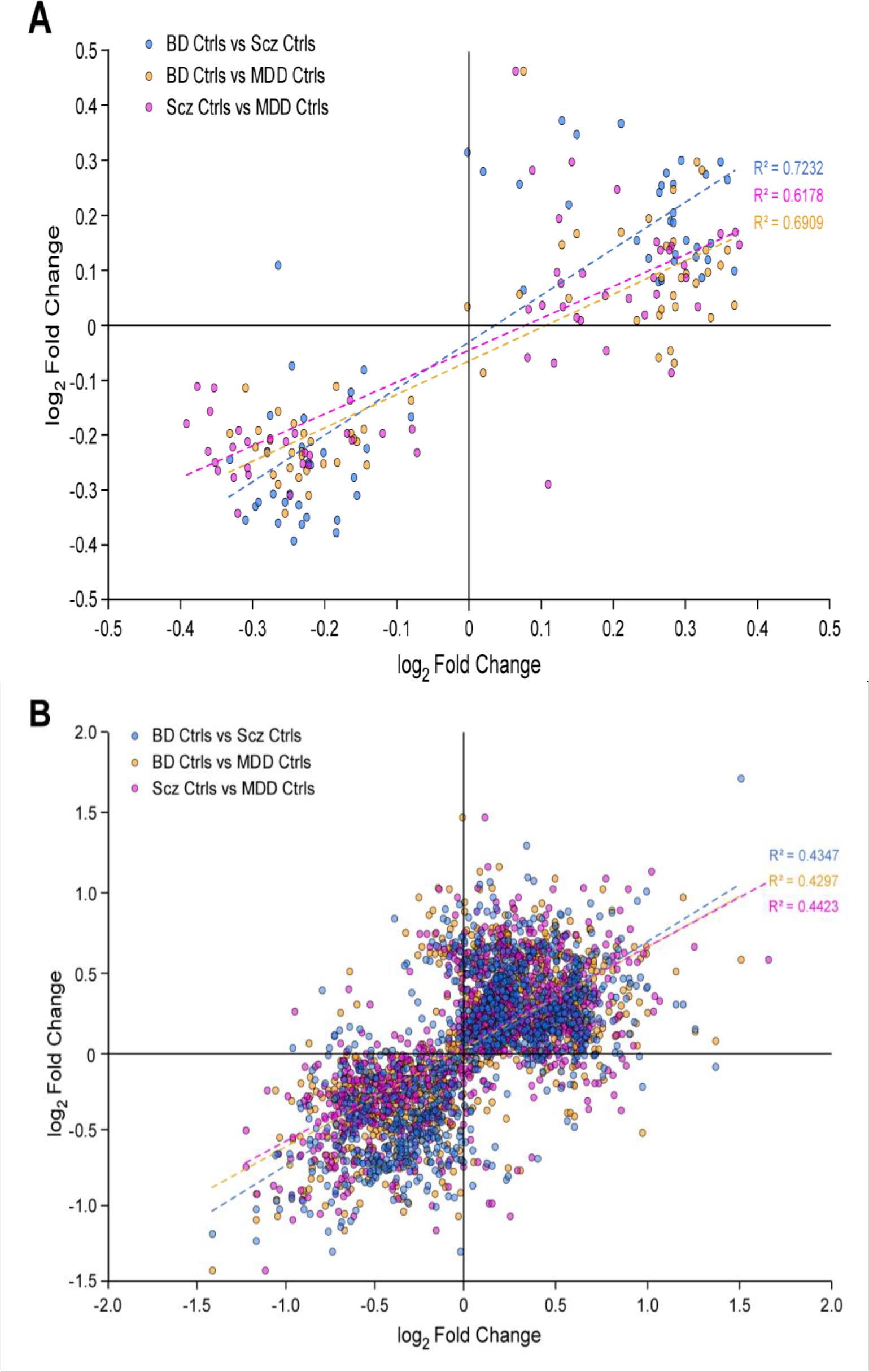
Correlation in case-control log_2_ fold-change values across diagnostic groups for a. genes and b. transcripts that were differentially expressed at FDR<5% in any cse-control comparison.

To confirm these results, the data were normalized, corrected for covariates, and analyzed using analysis of variance (ANOVA). Over 80% of the DE genes detected by DESeq2 were supported (data not shown).

#### Case-Case comparisons

In order to directly interrogate differential expression between diagnostic groups, we carried out analyses in all possible case-case comparisons (BD-SCZ, MDD-SCZ and MDD-BD). This approach revealed a greater diversity of DE genes and larger FC values than the case-control comparisons (Figure 1b). Seventy-three genes were DE in BD vs SCZ at FDR<5%, one of which, *HBG2*, was expressed in opposite directions in BD vs SCZ, compared to controls. At the same FDR 5% threshold, there were 9 genes that were DE in MDD vs SCZ and 5 genes that were DE in MDD vs. BD. The gene *IDUA* was expressed in opposite directions in each of these disorders, relative to controls. Supplementary tables 7, 8 and 9 show all genes DE at p<0.05 in each of the case-case comparisons.

#### Expression Quantitative Trait (eQTL) analyses

Genome wide association studies (GWAS) have identified numerous common variants associated with SCZ, BD, or MDD. In order to assess the effects of these variants on gene expression, we carried out an eQTL analysis with Matrix eQTL (see Methods). All samples were genotyped, imputed and filtered for common markers. This produced ~6M single nucleotide polymorphisms (SNPs) that were tested against expression of ~21K genes, resulting in ~92M eQTLs in cis (within 1MB upstream or downstream of a gene). Of these, ~290K eQTLs were associated with expression of 6,272 genes at FDR<5% (designated “eGenes,” Supplementary Table 10). We performed a similar analysis only in the Caucasian samples, resulting in ~99K cis-eQTLs for 4K genes at FDR<5%.

eGenes from sgACC were compared to the ~9K eGenes reported in DLPFC by the Common Mind Consortium^9^ (CMC-DLPFC) and to the ~4K eGenes reported by GTEx^16^ in anterior cingulate cortex (GTEx-ACC). About 50% of eGenes detected in sgACC overlapped with either CMC-DLPFC or GTEx-ACC (Supplementary Figure 5). eGene overlap with GTEx-ACC and CMC-DLPFC is presented in Supplementary Table 11. A total of 2,741 eGenes were detected only in sgACC, 881 of which were not found in the largest postmortem brain study to date^6^. Of these putative sgACC-specific eGenes, 521 had low expression in sgACC, with base mean < 100 counts (Supplementary Table 11), suggesting that they may have been missed by less deep RNA sequencing.

#### Summary-data-based Mendelian Randomization (SMR) analyses

In order to identify SNPs that were pleiotropically associated with both clinical diagnosis and gene expression in brain, we integrated published GWAS with brain eQTLs using SMR^17^. To increase power, we combined our eQTL results (Caucasian-only) with those detected in GTEx-ACC (~100 samples) and those found in CMC-DLPFC (~500 samples), thus boosting the total sample size to ~800.

As reported previously^18^, sample size was strongly predictive of the number of significant SMR “hits.” In SCZ, we detected 20 pleiotropic variants in sgACC at FDR<5%. This number increased to 36 in the combined sgACC and GTEx-ACC samples, and increased further to 133 when the CMC-DLPFC samples were added (Supplementary Table 12). Some of the genes reported here for the first time in association with a psychiatric GWAS locus include: *CORO7*, an F-actin regulator thought to play an essential role in maintenance of Golgi apparatus morphology; *TSHR*, which codes for the thyroid stimulating hormone receptor, previously reported to harbor rare, loss-of-function mutations in ADHD^19^; and *ORMDL3*, which plays a role in sphingolipid synthesis and regulation of Ca2+ levels in the endoplasmic reticulum^20^.

Integration of our eGene data with the latest BD GWAS results detected 4 pleiotropic variants, increasing to 16 when all brain samples were considered (Supplementary Table 13). This SMR analysis replicated 4 genes reported in previous SMR studies of BD ^6,21,22^: *DDHD2*, *MCHR1, FADS1*, and *PACS1.* The present study detected 5 novel BD eGenes, including: *ORMDL3*, noted above in relation to SCZ; *XPNPEP3*, an ezyme involved in glomerular filtration, protein processing, and alternative splicing; *PDF*, a mitochondrial protein involved in cell proliferation; and 2 non-coding RNA genes.

Similar analyses in MDD yielded 4 significant hits, but only in the combined dataset. (Supplementary Table 14). Exemplary genes include *DLST*, implicated in a recent anxiety GWAS^23^; and *B3GALTL*, mutations in which causes Peters-plus syndrome, characterized by developmental delay^24^.

### Transcript-level Analyses

#### Case-Control comparisons

RNAseq was carried out at high read depth in order to capture the highest number of detectable transcripts from the sgACC. Transcript-level comparisons between cases and controls showed 2-3 times more significant differences than the gene-level comparisons, despite a greater multiple-testing burden. At FDR<5% the number of DE transcripts was: 470 in SCZ, 380 in BD, and 286 in MDD, a gradient that is consistent with typical clinical severity across disorders (Figure 1c). Most (>90%) of the DE transcripts were predicted to be protein-coding (Supplementary Table 15). Many of the DE transcripts showed read counts <100, and were detected as DE in mental illness for the first time in this study (Supplementary Figure 6). All DE transcripts in SCZ, BD and MDD with p<0.05 are presented in Supplementary Tables 16-18.

In contrast to the gene level analyses, transcript-level analyses detected substantial diversity in DE transcripts across disorders. The mean overlap of DE transcripts across disorders was 18% (Figure 2b), and the correlation of FC values across diagnoses (linear regression R^2^) was 0.43-0.44% (Figure 3b). For some genes, the same transcript was DE in all 3 disorders (e.g., *NR4A1*-213, Figure 4), but there were several instances of alternative isoform usage, where distinct transcripts of the same gene were DE in different disorders (e.g., *HDAC11, NR4A1;* Figure 4).

**Figure 4:**
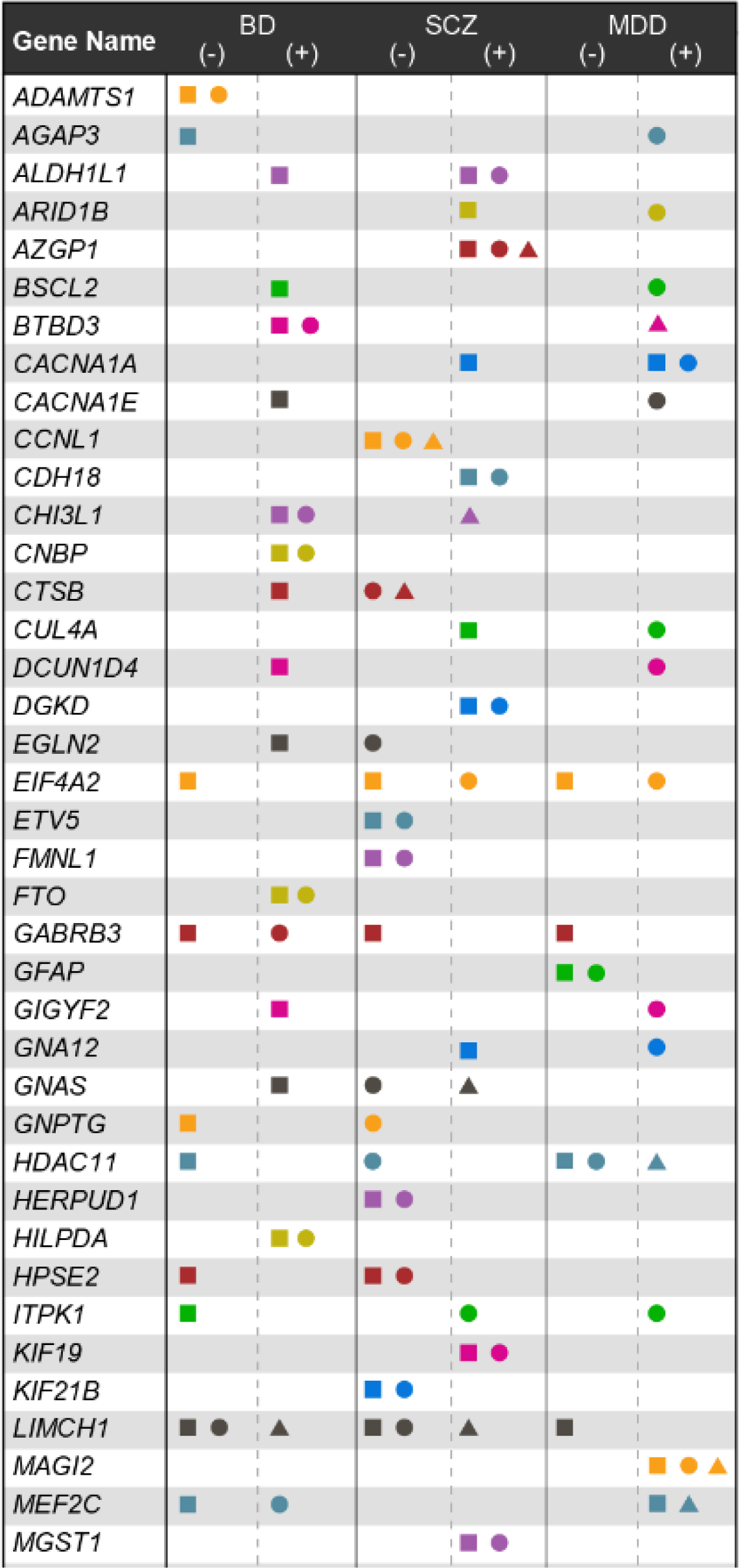
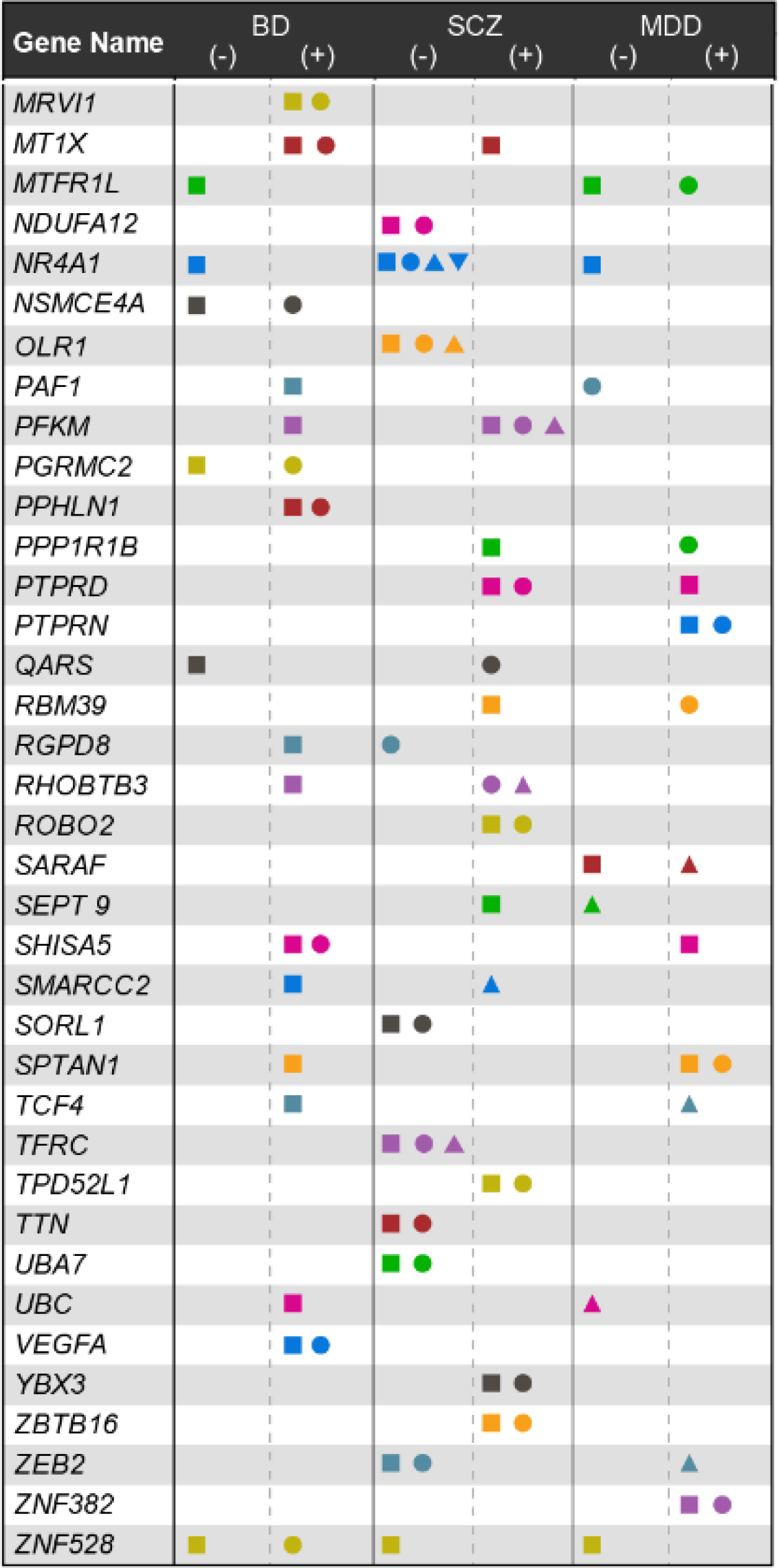
Genes with multiple differentially-expressed transcripts in bipolar disorder (BD), schizophrenia (SCZ), or major depressive disorder (MDD). Within each diagnosis column, downregulated transcripts are shown to the left, under “-“, upregulated transcripts are shown to the right, under “+”. Different colors indicate distinct genes; different shapes indicate distinct transcripts. Full gene and transcript names are shown in Supplementary tables 16-18.

A comparison of DE transcripts (FDR<5%) in SCZ and BD to those reported in the PsychENCODE sample^6^ showed small but significant overlaps (SCZ: hypergeometric p-value = 0.018; BD: hypergeometric p-value = 0.039), despite methodologic differences. Among transcripts previously reported as DE in PsychENCODE^6^, the present study replicated 22 transcripts in SCZ and two in BD (Supplementary Tables 16 and 17, respectively).

#### Case-Case comparisons

To further investigate the degree to which diagnoses could be differentiated at the transcript level, a series of case-case comparisons was performed, in which each psychiatric case group was compared to all the others. At FDR<5% we detected 466 transcripts that were DE between SCZ vs BD, 284 between SCZ vs MDD, and 264 between BD vs MDD (Figure 1d). DE transcripts with p<0.05 in any of the case-case comparisons are presented in Supplementary Tables 19-21.

While comparable to the case-control comparisons in number of detected DE transcripts, the case-case comparisons detected largely distinct sets of DE transcripts. For example, in the SCZ vs BD comparisons, DE transcripts included transcripts of: *TCF4*, *MBP*, and *LRRC4*. The SCZ vs MDD analysis detected DE transcripts in *CDK18*, *GNAS*, and *GRIK5*, among others (Supplementary Table 20). Transcripts DE in BD vs MDD included those within *SNAP91*, *MEF2C*, and transcripts of genes within established GWAS loci, including *SYNE1, MBP*, and *PBRM1*, among others (Supplementary Table 21).

Most of the DE transcripts detected in the case-case comparisons were expressed in opposite directions, in each case group, compared to controls. Opposing expression was found in 77% (SCZ vs BD), 83% (SCZ vs MDD), and 77% (BD vs MDD) of DE transcripts in the case-case comparisons (Figure 5a-5c; Supplementary Table 22-24). Since the absolute difference between case groups is larger for opposing transcripts (Figure 5d-5e), when power is limited such transcripts should be more easily detected in case-case than in case-control comparisons.

**Figure 5.**
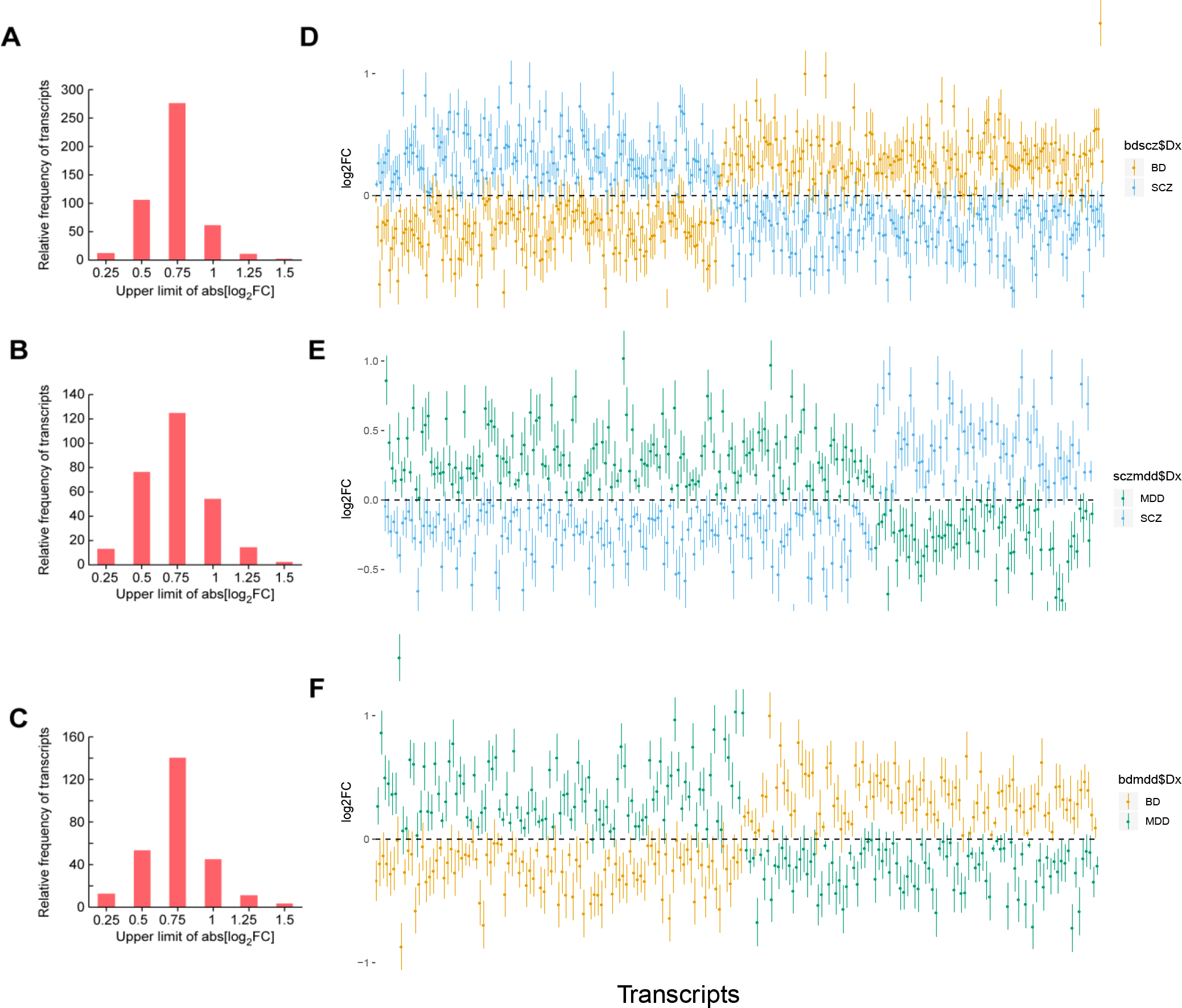
Transcript-level comparisons between case groups. Histograms of absolute log2 fold-change values are shown on the left; direction and magnitude (log_2_ fold-change) of expression in each case group relative to controls is shown on the right (for clarity, only transcripts expressed in opposite directions are shown, and x-axes are adjusted to the same length in each graph, although the number of transcripts differs). Schizophrenia (SCZ) vs bipolar disorder (BD) comparisons are shown in panels A. and D., SCZ vs major depressive disorder (MDD) comparisons are shown in A. and E., and BD vs MDD comparisons are shown in C. and F.

Comparisons of major mood disorders (BD and MDD) compared to SCZ found additional transcripts that were regulated in the same direction in the mood disorders and in the opposite direction in SCZ (Supplementary Table 25). Examples include transcripts of *HDAC10*, *SYNE1*, *MEF2C*, *SETD5*, and *TMEM132*. All of these genes have been implicated in mood disorders and/or SCZ by previous GWAS^21,25,26^.

#### eQTL analyses

At the transcript-level we detected ~200K eQTLs affecting expression of ~6K transcripts (FDR<5%) (Supplementary Table 26). These eQTLs significantly overlapped with transcripts that were DE in SCZ (hypergeometric p-value<10^−5^) or in BD (hypergeometric p-value<0.01), but not in MDD, suggesting that common alleles contribute to disorder-specific differential expression at the transcript level.

#### SMR analyses

To quantify the degree to which risk alleles contribute to transcript-level differences between disorders, SMR analysis was carried out with the transcript-eQTLs and summary results from recent GWAS. SNPs pleiotropically associated with both transcript expression and disease (FRD<5%) included 18 associated with SCZ and 4 associated with BD. Some of the significant SCZ loci include transcripts of *FAM134*, *KATNAL2*, and *ZNF738*. Signifcant BD loci include transcripts of *FADS1*, *XPNPEP3, MAP1LC3A*, and *ARPC3* (Supplementary Table 27).

### Validation of DE genes and transcripts

Validation of the RNAseq analyses was carried out in the same 185 samples using real time quantitative PCR (RT-qPCR) using 6 genes and transcripts with read counts over 500 (see Methods). All tested genes and transcripts showed the same direction of effect with respect to diagnosis as was detected in the RNAseq analyses. This result was significantly different from chance (binomial p-value of 0.016).

### Functional enrichment of DE genes and transcripts

Functional enrichment of analysis was carried out with DAVID.^27^ No significant functional enrichment was detected for any gene-level results, but DE transcripts did reveal some significant enrichment, consistent with the improved resolution reported above for transcript-level comparisons. Genes harboring transcripts that were DE at FDR<5% in any of the case-control comparisons were significantly enriched for functional annotations related to synapse and antigen processing (Supplementary Table 28), largely mirroring the enrichment detected in the SCZ vs Ctrls comparison (Supplementary Table 29). In case-case comparisons (Supplementary Tables 30), SCZ vs BD revealed significant enrichment for functional annotations related to muscle or motor proteins, while SCZ vs MDD highlighted spectrin repeats and cell membrane. No significant enrichment was found for the small number of DE transcripts detected in the BD vs MDD comparison.

### Effects of Common Variants on Transcript Abundance

In addition to changes in gene or transcript-level expression, SNPs can modify the transcriptome by driving shifts in the relative abundance of transcripts within a gene^28^. Such SNPs are known as sQTLs, due to their putative effect on alternative splicing^29^. Accordingly, we assessed the relationship between common variants and relative abundances of all detected transcripts in each of the ~21K genes included this study. This analysis detected ~90K sQTLs associated with transcript abundance of ~5K genes at p-value<0.05 (Supplementary Table 31).

A similar analysis restricted to the Caucasian samples detected ~75K sQTLs in ~4K genes at p-value<0.05 (Supplementary Table 32). As expected, most (63%) transcripts associated with one or more sQTLs were predicted to result from classical alternative splicing events, including exon skipping, intron retention, alternative donor, or alternative acceptor sites (Supplementary Figure 7). These genes overlapped significantly (hypergeometric p < 1.11e-16) with those identified by different methods in the CMC data^30^. About 20% of these sQTLs are intronic variants located within 1 KB of a known splice site (Supplementary Table 33).

Genes harboring one or more sQTLs significantly overlapped wiht genes harboring DE transcripts from the case-control (n=368; OR=2.2, hypergeom p<10^−16^) and the case-case (n=335; OR=2.5, hypergeom p<10^−16^) comparisons. No significant functional enrichment was detected for the overlapping genes from the case-control analyses, but case-case overlaps were significantly (FDR<5%) enriched for synapse, post-synaptic density, cell junction, and spectrins. Box plots of genes harboring sQTLs and DE transcripts are shown in Supplementary Figure 9.

#### sQTL enrichment in regulatory regions

Regulatory annotation of sQTLs was peformed with the Encode regulatory database as implemented in SNPnexus^31^. The most common annotations included the histone marks H3K36me3, H3K4me1, and H3K79me2, DNAse1 hypersensitive sites, and predicted binding sites for the transcription factor, CTCF (Figure 6; Supplementary Table 34). These results are consistent with a role for sQTLs in alternative splicing^32^.

**Figure 6:**
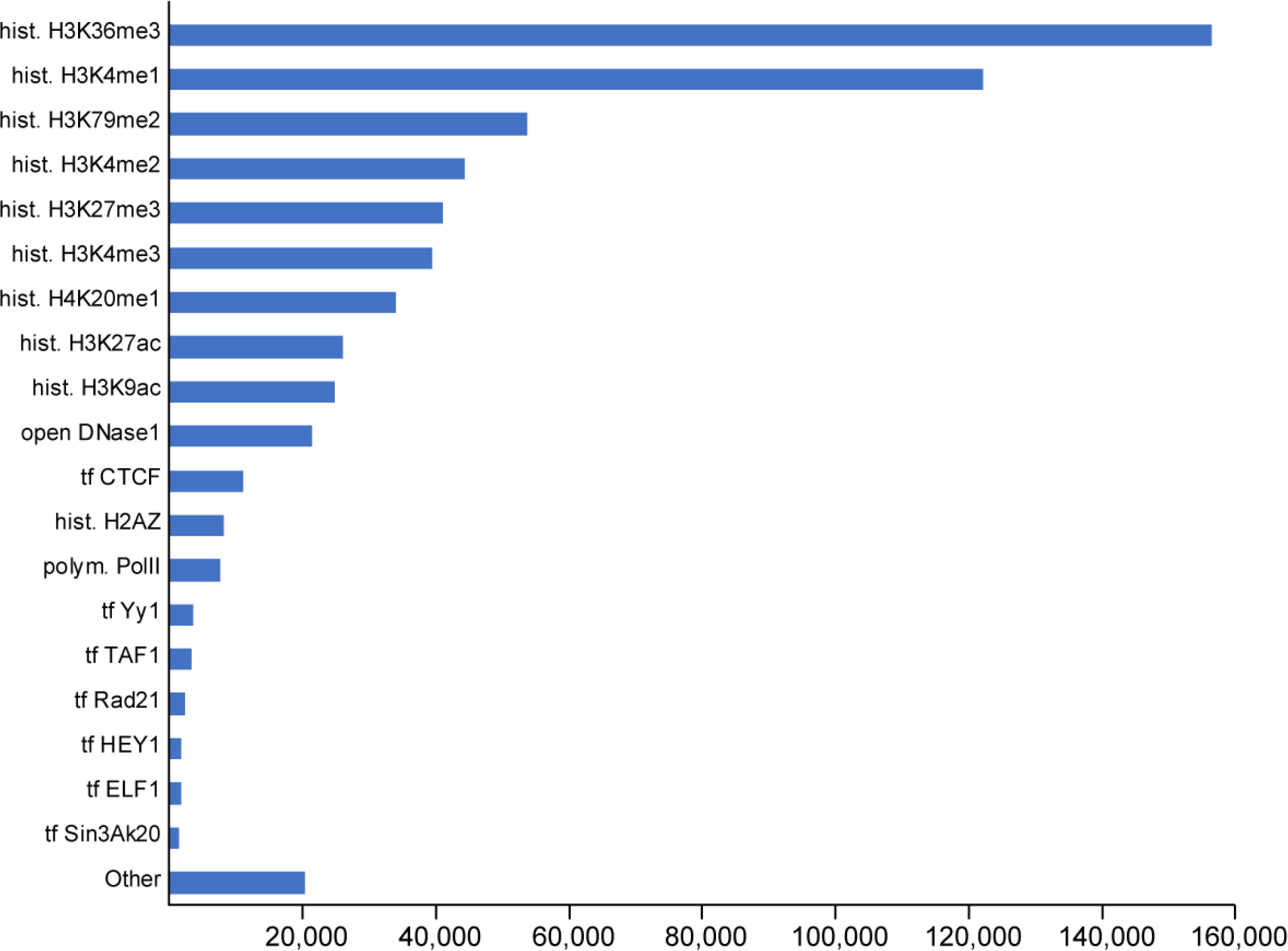
Regulatory annotation of sQTLs. ENCODE annotation terms are shown on the y-axis, counts of identified sQTLs (p<0.05) are shown on the x-axis.

#### sQTLs within GWAS loci

As observed with eQTLs, some sQTLs may be pleiotropically associated with disease risk. To investigate this, we searched for SNPs that were identified as risk alleles by GWAS and as sQTLs in the present study. We used only those sQTLs that were detected in sgACC from self-reported Caucasian brain donors, since existing GWAS are mainly based on samples of European ancestry,.

Overall, about 10-25% of GWAS loci we investigated harbored one or more significant sQTLs. In SCZ, of 430 genome-wide significant GWAS SNPs^25^, 56 were identified as an sQTL in the present study, implicating 44 genes. Of these, 15 different genes are linked here for the first time to functional variation within a SCZ GWAS locus (Supplementary Table 35). In BD, of 329 genome-wide significant GWAS SNPs^21^, 47 were identified as sQTLs (Supplementary Table 36. Some of the novel implicated genes include *ATP11B, NCAM1*, *SLC12A9*, and *SLC4A10*. In MDD, 10 out of 44 GWAS loci^26^ were found to harbor sQTLs, implicating the genes *BAG6*, *KCNQ5* and *ERBB4*, among others (Supplementary Table 37).

#### Relative contributions of eQTLs and sQTLs to disease heritability

Previous studies have demonstrated a substantial contribution of eQTLs to heritability for a variety of common disoders^33^, but few studies have investigated the specific contribution of sQTLs. Thus we estimated the proportion of heritability that could be explained by sQTLs, relative to eQTLs, in some major mental illnesess.

Both eQTLs and sQTLs accounted for disproportionate fractions of heritability in SCZ, BD, and MDD (Table 3). However, sQTLs, which comprised only 1% of all SNPs, explained 16-21% of the heritability, a 14-18 fold enrichment (p<0.01). sQTLs showed even greater enrichment of heritability for some other major psychiatric disorders, but not all. sQTLs explained 29% of the heritability of autism spectrum disorders (25-fold enrichment, p<0.001) and 41% of the heritability of attention deficit hyperactivity disorder (36.6-fold enrichment, p<0.001), but showed no significant enrichment for Alzheimer Disease or anxiety disorders.

**Table2:** Heritability of psychiatric disorders, partitioned across sQTLs and eQTLs

| Diagnosis | N SNPs | Observed H2 (se) | h2 using sQTL+eQTL (StdErr) | Proportion sQTLs | sQTL h2 (se) | sQTL Enrichment | sQTL Enrichment se | sQTL Enrichment p | Proportion eQTLs | eQTL h2 (se) | eQTL Enrichment | eQTL Enrichment se | eQTL Enrichment p |
| --- | --- | --- | --- | --- | --- | --- | --- | --- | --- | --- | --- | --- | --- |
| Schizophrenia | 1,069,224 | 0.4 (0.015) | 0.12 (0.009) | 0.011 | 0.20 (0.03) | 18 | 2.8 | 6.70E-09 | 0.047 | 0.80 (0.03) | 16 | 0.66 | 1.36E-21 |
| Bipolar disorder | 962,367 | 0.3 (0.018) | 0.07 (0.008) | 0.011 | 0.16 (0.04) | 14.4 | 3.5 | 1.23E-04 | 0.047 | 0.84 (0.04) | 17.6 | 0.84 | 2.07E-13 |
| Major depression | 1,060,414 | 0.05 (0.003) | 0.008 (0.001) | 0.011 | 0.21 (0.071) | 18 | 6.3 | 7.57E-03 | 0.047 | 0.80 (0.071) | 16.7 | 1.5 | 4.80E-08 |
| Autism spectrum disorders | 1,049,026 | 0.18 (0.017) | 0.03 (0.005) | 0.011 | 0.29 (0.08) | 25.3 | 7 | 7.62E-04 | 0.047 | 0.71 (0.08) | 15 | 1.65 | 3.19E-05 |
| Attention deficit hyperactivity disorder | 999,575 | 0.21 (0.015) | 0.026 (0.006) | 0.011 | 0.41 (0.11) | 36.6 | 10.1 | 6.81E-04 | 0.047 | 0.59 (0.11) | 12.3 | 2.4 | 1.00E-02 |
| Alzheimer disease | 1,035,603 | 0.046 (0.009) | 0.017 (0.003) | 0.011 | 0.13 (0.08) | 11.4 | 7.4 | ns | 0.047 | 0.87 (0.08) | 18.3 | 1.8 | 1.66E-06 |
| Anxiety | 897,467 | 0.06 (0.033) | - | 0.011 | - | - | - | ns | 0.047 | - | - | - | - |

\* Excludes eQTLs that are also sQTLs

## DISCUSSION

We have employed deep transcriptome sequencing in post-mortem brain tissue to address a central question in the pathophysiology of mental illness: How do diagnostic differences in onset, symptoms, and treatment response arise despite substantial overlaps of genetic risk factors across disorders? The findings show that subtle differences in gene expression observed between broad diagnostic classes of mental illness actually reflect more pronounced and diagnosis-specific changes at the transcript level. These changes often involve alternative isoform usage, are most evident in case-case comparisons, and are influenced by common genetic variants that also play a disproportionately large role in the inherited risk of mental illness.

These findings should be considered in light of several limitations. While this was one of the larger RNA sequencing studies performed to date in human brain tissue, sample size was limited, with less than 100 individuals in each diagnostic category. This limits power to detect small changes in expression. This limitation was offset by complementing the typical case-control comparisons with case-case comparisons between distinct psychiatric diagnoses. This approach should be more powerful in the face of opposing expression changes between case groups, which we detected for hundreds of genes and transcripts. As in all human post-mortem studies, diverse factors effect post-mortem expression, not all of which can be measured or controlled, increasing noise and introducing the risk for bias. We attempted to address this by stringent quality control, careful adjustment for key covariates, and confirmation of top findings by use of alternative approaches. Also inherent in human post-mortem samples is the inability to draw confident distinctions between expression changes that cause disease and those that result from disease or its treatment. The kind of integrative genomic approaches we employed, which draw causal inferences on the basis of published GWAS^17^, partially address this issue but still lack power to distinguish between correlated causal and non-causal events. This study focused on a single brain region, albeit one that has been little studied, so cannot shed light on transcriptional differences between brain regions that may be important in disease. While we employed very deep RNA-sequencing to detect rare transcripts and better identify alternative splicing, the data still contain relatively few sequencing reads that uniquely map to splice junctions, limiting the confidence with which transcript-level differences can be ascribed to differences in alternative splicing rates and mechanisms. Finally, the sequencing was performed on bulk tissue. This means that the results lack cellular resolution, but also that they comprise a much larger portion of the brain transcriptome than can usually be achieved with current single cell technologies.

Several large postmortem brain gene expression studies of mental illness have been published recently, some with larger samples than were employed here^6,9,34^. The present study complements those larger studies in several ways. None of those studies evaluated the subgenual anterior cingulate cortex (sgACC), a key component of limbic circuits that has been implicated in reward, impulse control, and emotion regulation^5^, as well as in major mood disorders.^7,8, 35–37^ To date, genome-wide expression studies in ACC or sgACC have been carried out in fewer than 50 samples^38–40^. Some of the genes and transcripts whose expression we found to be associated with GWAS hits were detected only in ACC, consistent with the idea that common risk alleles may regulate expression in a tissue-specific manner^41^. None of the previous studies employed RNA-sequencing at a depth comparable to the 270 million reads per sample used here, which enabled us to capture at least 10 reads per sample in more than 21,000 genes and 85,000 transcripts. This was particularly beneficial in transcript-level analyses, which found numerous DE transcripts, shifts in relative transcript abundance that were associated with numerous common variants (sQTLs), and strong evidence that such variants play a disproportionately large role in the heritability of SCZ, BD, and MDD. The present study was also unique in its use of case-case comparisons, which revealed unexpectedly large numbers of genes and transcripts that were DE between case groups. Many DE genes and transcripts showed opposite directions of expression between cases, when compared to controls, suggesting one important way in which genetically correlated disorders can express contrasting transcriptomes that could reflect diagnostic differences in onset, symptoms, or treatment response.

Most of the transcripts whose expression was associated with common genetic variants in this study are predicted to arise from classical alternative splicing mechansims. These findings are consistent with previous results suggesting that alternative splicing plays an important role in differential transcriptome expression in major psychiatric and other disorders and that it is mediated, at least in part, by some of the same genetic variants identified by GWAS^32^. A secondary analysis of RNA-sequencing data from psychiatrically unaffected individuals drawn from the CMC dataset found evidence of alternative splicing associated with common genetic variants, some of which overlapped with known SCZ risk loci, but did not assess for differential expression in individuals diagnosed with SCZ^30^. A recent study of post-mortem human cortex at 85 million reads per sample detected shifting isoform usage across pre-and post-natal life and a number of transcripts that were DE in SCZ^41^. Our results support those previous findings and demonstrate that common genetic variants associated with relative transcript abundance (sQTLs) account for disproportionate fractions of disorder-specific heritability.

Taken together, these findings suggest that inherited genetic risk factors shape the brain transcriptome and contribute to diagnostic differences between broad classes of mental illness. This may be one important mechanism by which diagnostically distinctive patterns of transcript expression arise despite the known correlations in inherited genetic risk factors across major psychiatric disorders. Despite the growing recognition of the importance and complexity of alternative splicing, its role in the development of neuropsychiatric diseases is still poorly understood. Much future work is needed to characterize the functions of alternative transcripts, the timing of alternative isoform usage during disease course, and the potential impact that therapeutic agents may have on the expression or function of particular isoforms in specific brain regions and cell types.

## METHODS

See Methods document

### Data availability

The raw data can be downloaded from dbGAP at https://www.ncbi.nlm.nih.gov/projects/gap/cgi-bin/study.cgi?study_id=phs000979.v2.p2

### Tissue availability

Tissue samples are available to qualified scientists upon review by the NIMH Human Brain Collection Core oversight committee. Information is available at https://www.nimh.nih.gov/research/research-conducted-at-nimh/research-areas/research-support-services/hbcc/distribution-of-resources.shtml

## ACKNOWLEDGEMENTS

This study was supported in part by the NIMH Intramural Research Program (ZIC MH002903-12, ZIAMH002810). We would like to thank the patients and their families for the donation of brain tissue for research. Data was analyzed on the high-performance Biowulf cluster at NIH. Library preparation and RNA sequencing was performed at the NIH Sequencing Center,

## Author Contributions

NA, FJM conceived and planned the project. BTH, JA, SM, CHR, and BKL performed neuropathology and case selection. NF, JC, BE, and QX performed the experiments. NA, KJ, KA and RK performed data analyses. NA and FJM wrote the manuscript and SM, SDW, PKA, and BKL edited the manuscript. All coauthors had the opportunity to review and approve the manuscript before submission.

## METHODS

### Samples

All the 200 post-mortem brain samples (61 controls; 39 bipolar disorder; 46 schizophrenia; 54 major depression) were collected by the Section on Neuropathology of the Clinical Brain Disorders Branch at the National Institute of Mental Health (NIMH) under protocol# 90-M-0142^1^. All samples were dissected from frozen coronal slabs cut at autopsy. The portion of anterior cingulate cortex underlying the genu of the corpus callosum (generally including Brodmann areas 25, 32 and 24) was targeted. Dissections were done by dental drill cleaned between specimens (H2O/Bleach/H2O). Specimens were laid out on a cutting board placed on dry ice and kept frozen during dissections. Following dissection, total RNA was extracted (extraction batch in Supplementary Table 1) from 50mg of pulverized subgenual anterior cingulate cortex region. Samples with RIN <6 were excluded from sequencing.

### RNA sequencing

Stranded RNA-Seq libraries were constructed from 1 µg total RNA after rRNA depletion using Ribo-Zero GOLD (Illumina Inc, San Diego, CA, USA). The library batch per sample is presented in Supplementary Table1. The Illumina TruSeq Stranded Total RNA Sample Prep Kit was used according to manufacturer’s instructions except where noted. Amplification was performed using 10 cycles which was optimized for the input amount and to minimize the chance of over-amplification. Libraries were pooled together for sequencing in equimolar amounts. The pooled libraries were sequenced on a HiSeq 2500 using version 4 chemistry. The data was processed using RTA version 1.18.64 and Casava 1.8.2. RNA sequencing was performed at National Institute of Health Intramural Sequencing Center (NISC). Stranded paired-end sequencing with read length of 125bp were generated. We had an average of 137 million reads per sample totaling to ~54 billion reads for all samples.

### Mapping and counting

The reads were trimmed using Trimmomatic with default parameters^2^ (http://www.usadellab.org/cms/?page=trimmomatic). The trimmed reads were then mapped to the human genome (Ensembl GRCh38.87) using Hisat2^3^ (https://ccb.jhu.edu/software/hisat2/index.shtml) with default parameters. Known genes and transcripts belonging to autosomes and pseudo-autosomal regions (PAR) were included in the analysis. Gene and transcript counts were obtained using StringTie^3^ software (http://ccb.jhu.edu/software/stringtie/index.shtml?t=manual.

### Sample and Gene selection

Pre-noise filtered quantile (Log_2_(FPKM+2)) expression values for all 200 samples were used to calculate Pearson correlations between each possible pair of samples within a diagnostic category. The average correlation for each sample within a diagnostic category was obtained, providing a single estimate of relatedness of that sample to all others in its diagnostic category. We then tested for deviance of each sample to the rest of their diagnostic group by a z-score test, choosing a conservative threshold of p<0.1 to define individual samples as outliers. By this approach, 15 samples were identified and removed as outliers (Supplementary Figure 10). Lowess modeling was performed by disease class to remove noise-biased expression, thus resulting in ~21K genes (Supplementary Figure 11).

### Covariates

We tested for a total of 23 known covariates: RIN, age, gender, reported race, pH, RNA extraction batch, library batch, postmortem interval (PMI), PMI confidence, GC percent, manner of death, smoking status, source location, height, and weight, along with available data on alcohol, anti-depressants, mood stabilizers, benzodiazepines, nicotine, THC, cocaine, and opiates. Univariate and step-wise AIC regression analysis was performed on all the covariates. Any covariates that were significant after the multiple testing were included in the downstream analysis.

### Gene-and transcript-level analysis

We normalized the raw counts using conditional quantile normalization^4^ (CQN) (https://bioconductor.org/packages/release/bioc/html/cqn.html) which corrects for library size, gene length and GC content. These normalized counts were then imported to DESeq2^5^ (https://bioconductor.org/packages/release/bioc/html/DESeq2.html) and transformed to variance stabilized data (VSD). Principal component analysis (PCA) using VSD data resulted in 10 PCAs all of which together explain >97% of variance (each PCA >= 0.3%). Each of the 10 PCAs were linearly regressed against the 23 known covariates. Only 3 covariates – RIN, race and GC percent were significant after correcting ted for multiple tests, so these were included in the downstream analysis. A total of 185 samples (55 controls, 35 bipolar disorder, 44 schizophrenia, 51 major depression) were included in the analysis after removing 15 outliers. A total of 21,228 genes and 85,295 transcripts were included in the analysis. DESeq2 along with lfcShrink option was used for differential expression analysis between each diagnostic group versus the controls. The p-values were corrected using FDRtool^6^ software in R.

### Analysis of variance (ANOVA) analysis

DE genes (p<0.05) in each of the disorder were validated by ANOVA after correcting for the covariates. Dunnet’s multiple comparisons tests were used as follow up tests to ANOVA.

### DAVID Functional annotation

Functional annotation was performed in DAVID which is a database of annotation, visualization and integrated discovery^7^ (https://david.ncifcrf.gov/content.jsp?file=citation.htm). All 85,295 transcripts that met criteria for inclusion in the DE analysis were used as background genes.

### Genotype data and Imputation

Samples were genotyped on Illumina HumanHap650Y, Omni1-quad, Omni5v1.0 and Omni5v1.1 arrays (Illumina Inc, San Diego, CA, USA). Markers that have minor allele frequency (MAF) <1%, genotype missing rate >5% and Hardy-Weingberg Equilibrium (HWE) p<10^−4^ were excluded. All individuals had >95% genotype rate (--mind 0.05). Common markers across the multiple arrays were extracted and files were merged using PLINK2.0 (https://www.cog-genomics.org/plink/2.0/). All the samples were imputed against the Haplotype Reference Consortium (HRC release 1.1 panel) and pre-phasing was done using EAGLE2 on Sanger imputation server^8^ (https://imputation.sanger.ac.uk). After imputation any markers with R^2^ < 50%, MAF < 5%, and multi-allelic variants were excluded. All the markers were converted from hg19 to GRCh38 positions.

### Expression quantitative trait loci (eQTL) analysis

Matrix eQTL software^9^ was used to identify the eQTLs that affect the gene/transcript expression (http://www.bios.unc.edu/research/genomic_software/Matrix_eQTL/). Variants within 1MB of a gene were classified as “cis”. All default parameters were used. In the transcript-level analysis, transcripts with base Mean<100 were excluded due to computational problems.

### eQTL comparison with CMC and GTExACC

SNPs in sgACC were mapped to hg38 while the Common Mind Consortium (CMC) and GTExACC were mapped to hg19. For comparison purposes we used SNPs with rs numbers rather than position. The Common Mind Consortium eQTL data was obtained through Synapse (https://www.synapse.org) and the GTExACC data was downloaded from the GTEx website (https://gtexportal.org/home/datasets)

### Splicing quantitative trait loci (sQTL) analysis

Variants associated with alternative splicing were identified by sQTLseekeR^10^, a R package, with default parameters (https://www.nature.com/articles/ncomms5698). The sQTLs were characterized using AStalavista^11^ software (http://genome.crg.es/astalavista/). Functional annotation of the sQTLs was performed using SNPnexus^12^ (http://www.snp-nexus.org)

### Publicly available datasets

The genome-wide association study (GWAS) results were obtained from PGC for Schizophrenia (http://www.med.unc.edu/pgc/results-and-downloads), Bipolar Disorder GWAS dataset and Major Depression GWAS dataset were obtained from PGC^13^ and UKbiobank^14^ (early access upon request). GWAS data for Autism Spectrum Disorders (ASD), Attention-Deficit/Hyperactivity Disorder (ADHD) and Anxiety were downloaded from PGC (https://www.med.unc.edu/pgc/results-and-downloads). The Alzheimers GWAS data was download from IGAP (http://web.pasteur-lille.fr/en/recherche/u744/igap/igap_download.php). The Common Mind Consortium (CMC) eQTL dataset for dorso-lateral prefrontal cortex region (DFPLC) was downloaded via synapse after data access agreement was approved (https://www.synapse.org). The GTEx eQTL data for anterior cingulate region (ACC) was obtained from GTEx portal (https://GTExportal.org/home/datasets).

### SMR analysis

The pleiotrophic variants and their corresponding genes were identified by analyzing the eQTL data and GWAS data in SMR software^15^ (https://cnsgenomics.com/software/smr/#Overview). Due to the complexity of the MHC region on chr6 (26MB-34MB), this region was excluded in the SMR analysis. All default parameters were used.

### Enrichment analysis

For Schizophrenia, GWAS summary results, including index and credible SNPs, were downloaded from the PGC website (https://www.med.unc.edu/pgc/results-and-downloads). For Bipolar disorder and Major depression we took GWAS hits (SNPs with trait association p-values<10^−6^) from the public summary statistics noted above and used PLINK^16^ (http://zzz.bwh.harvard.edu/plink/) to extract all SNPs in linkage disequilibrium (r^2^>0.6) with the GWAS hits (“LD friends”). We also used PLINK to calculate linkage disequilibrium values among the index SNPs, LD friends, and sQTLs (p<0.05). We then extracted sQTL-GWAS SNP pairs with r^2^>0.6. The pairs were then pruned to get independent loci using the “indep-pairwise” option in PLINK (--indep-pairwise 50 5 0.5). The 1000Genomes European data^17^ was used as reference (http://www.internationalgenome.org/data).

### Heritability estimate

Heritability estimates were performed with LDSC software^18^ (https://github.com/bulik/ldsc), using the GWAS summary statistics for each disorder. To perform the partitioned heritability analysis for sQTLs and eQTLs we created a custom annotation file according to the instructions given in LDSC software (https://github.com/bulik/ldsc/wiki/LD-Score-Estimation-Tutorial), designating each SNP as an sQTL or an eQTL that was not also an sQTL. This annotation file was then used to partition the heritability between sQTLs and eQTLs in each disorder.

### qPCR validation

Some genes and transcripts with expression that was high enough to produce reliable results by qPCR (baseMean > 500) and that were identified as DE in the DESeq2 analysis of the RNA-seq data were selected for validation. At the gene-level, we selected *DUSP1* and *NR4A2*, both of which were DE in BD, and *HIGD2A*, which was DE in BD and MDD. At the transcript level, we selected transcripts of *ARID5B* (ENST00000279873), *NT5DC3* (ENST00000392876) and *SF3A1* (ENST00000215793) that were DE in SCZ. Taqman probes were ordered from Applied Biosystems (Thermo Fisher Scientific, 168 Third Avenue, Waltham, MA, USA). Catalog numbers are Hs00610256_g1, Hs00431157_g1, Hs01117527_g1, Hs01381961_m1, Hs00213132_m1 and Hs01066327_m1. Raw cycle counts were normalized against Actin, *B2M*, and *GUSB*, and delta CT values were regressed against diagnosis, with RIN, race, and GC percent as covariates.

