## Supplementary Figures for "Deep transcriptome sequencing of subgenual anterior cingulate cortex reveals disorder-specific expression changes in major psychiatric disorders"

Supplementary Figure1: Analysis pipeline of subgenual anterior cingulate cortex (sgACC) transcripts

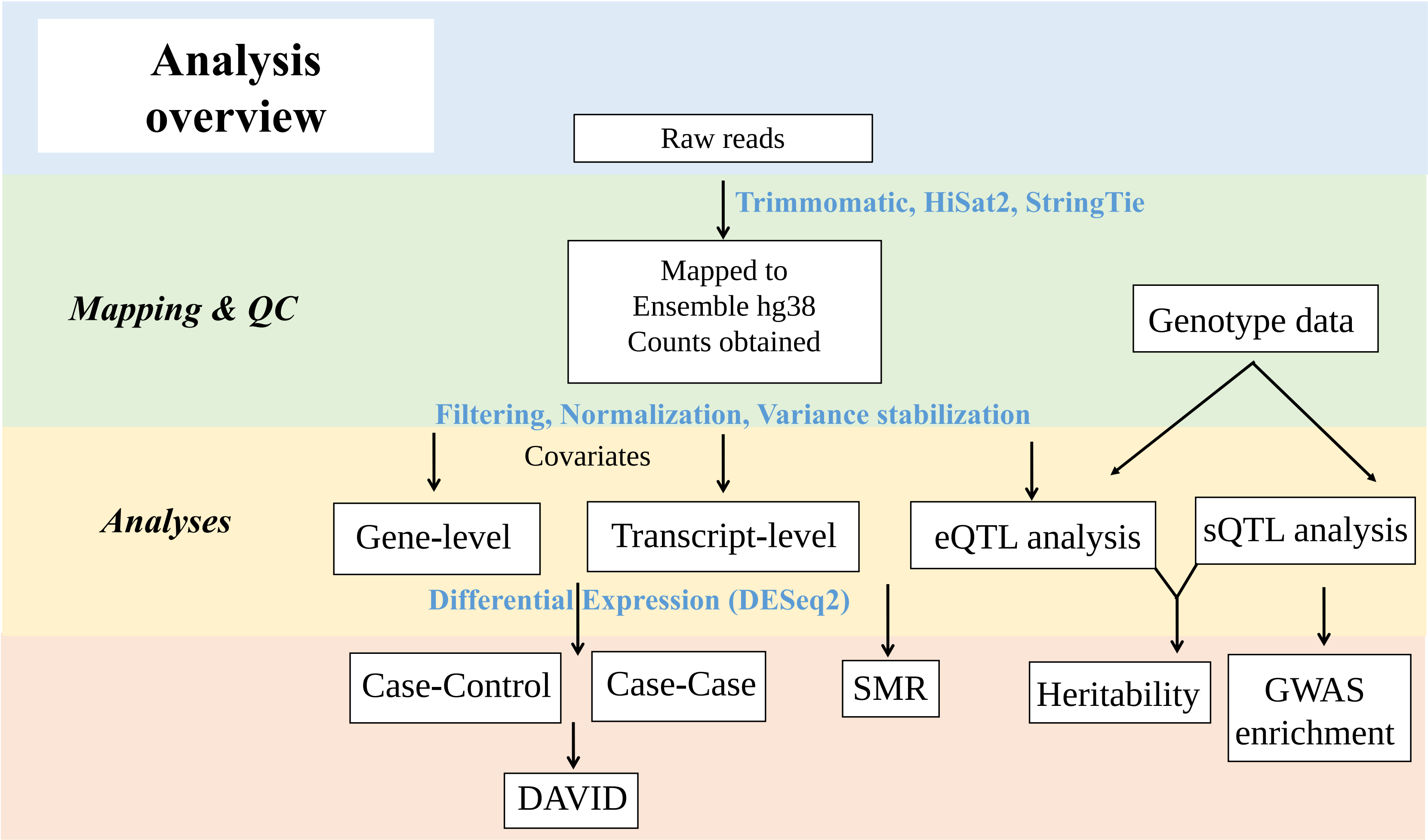

Supplementary Figure 2: Read count distribution of ~85K transcripts included in the study. The small number of transcripts with read counts >1,000,000 are not visible. Labels indicate upper bound of read counts.

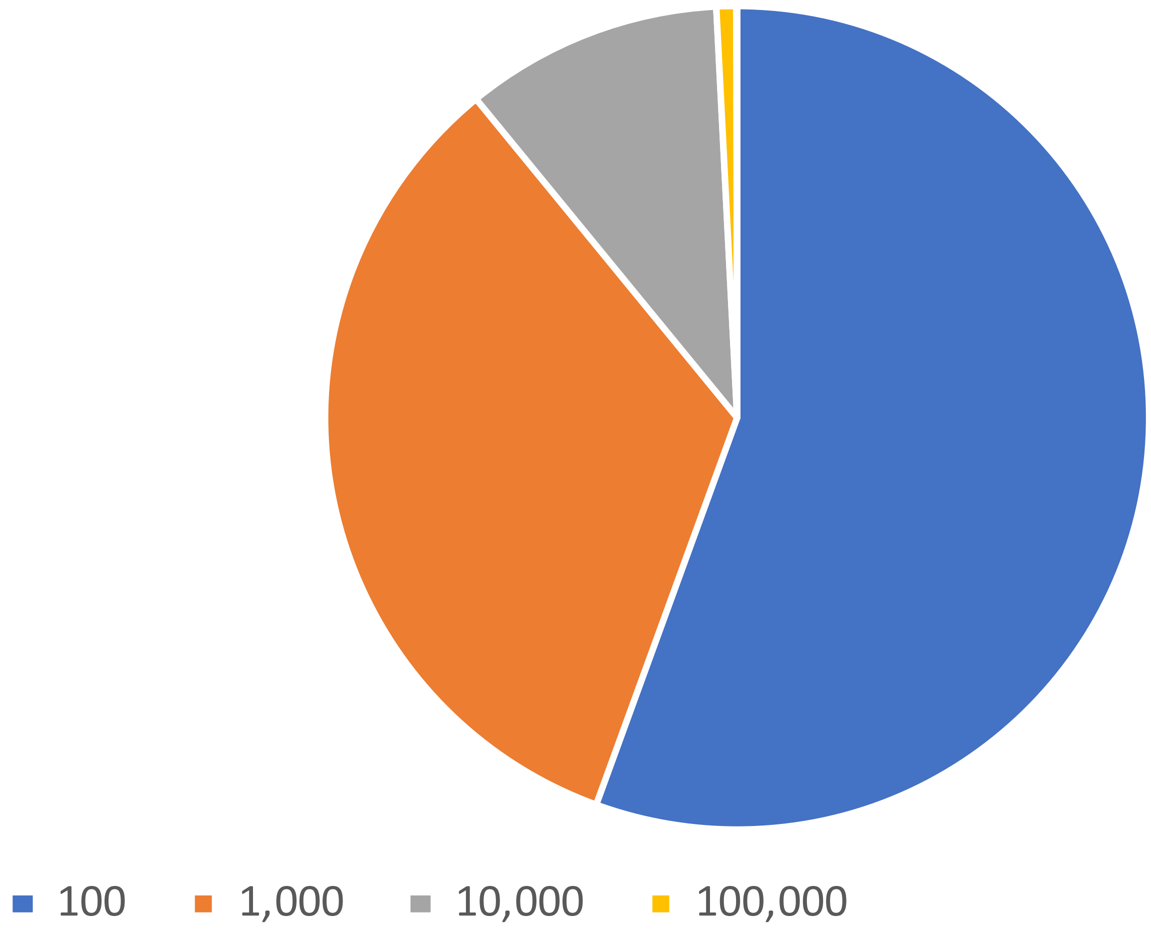

Supplementary Figure 3: QQ-plots depicting the observed (orange) and expected (blue) distribution of -log(p-values) for gene-level case-control differential expression results.

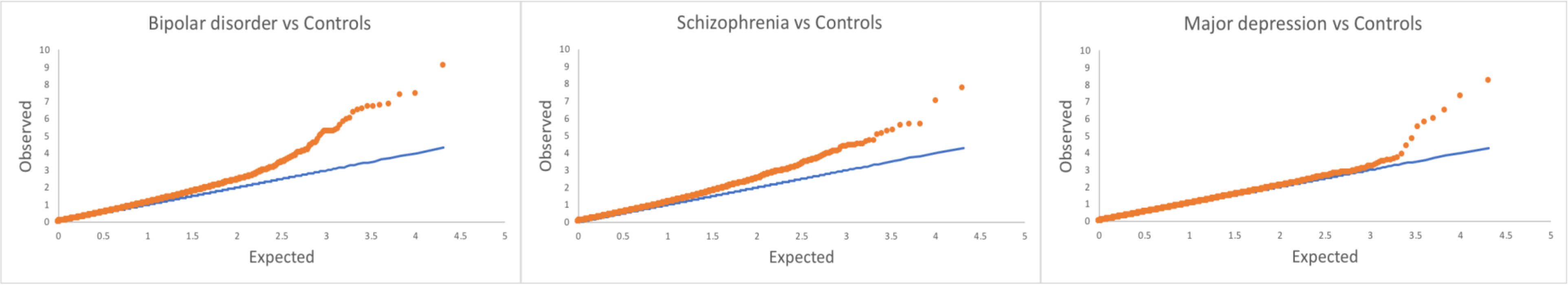

Supplementary Figure 4a: Gene-level: Volcano plots of all genes in case-case comparisons. Genes with FDR<5% are highlighted in red.

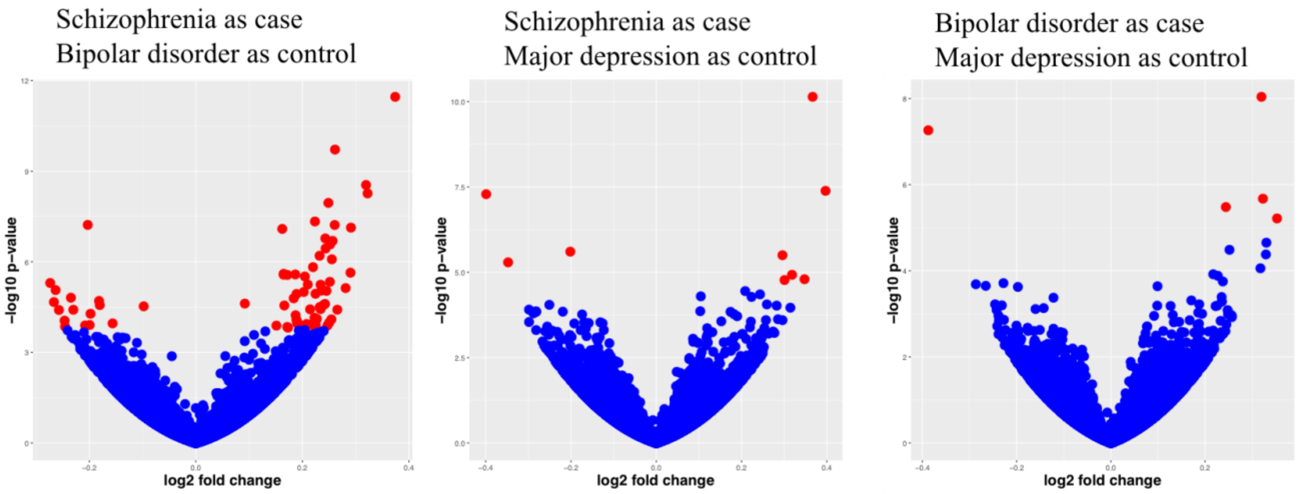


Supplementary Figure 4b: Transcript-level: Volcano plots for all transcripts in case-case comparisons. Red dots indicate genes with FDR<5%

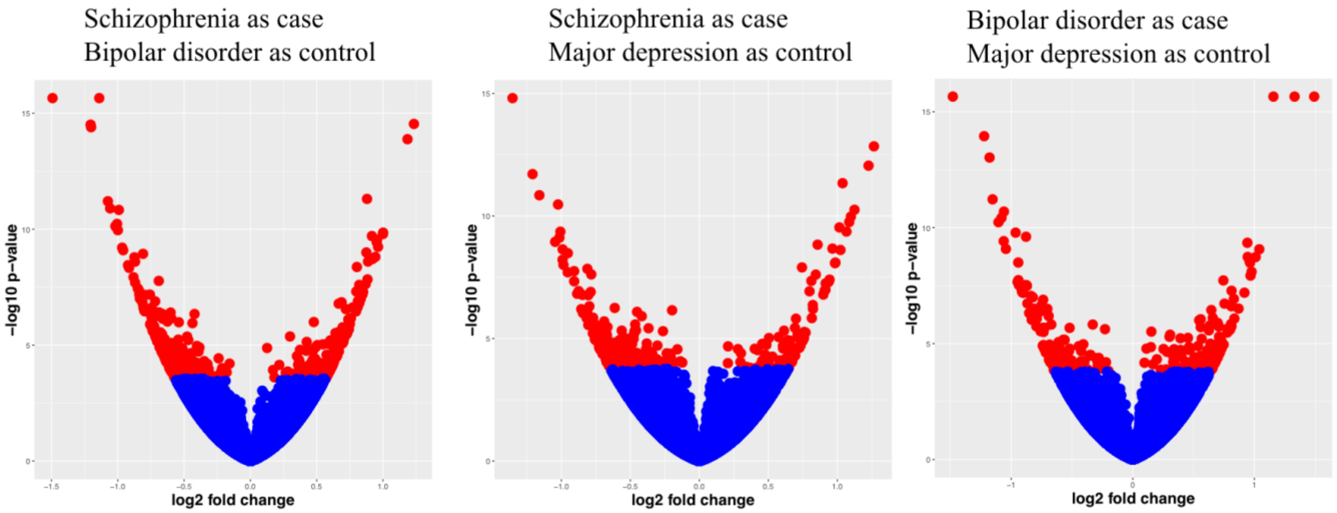

Supplementary Figure 5: eGenes (FDR<5%) overlap between subgenual anterior cingulate cortex (sgACC), Common Mind Consortium (CMC) and GTex anterior cingulate cortex (GTexACC).

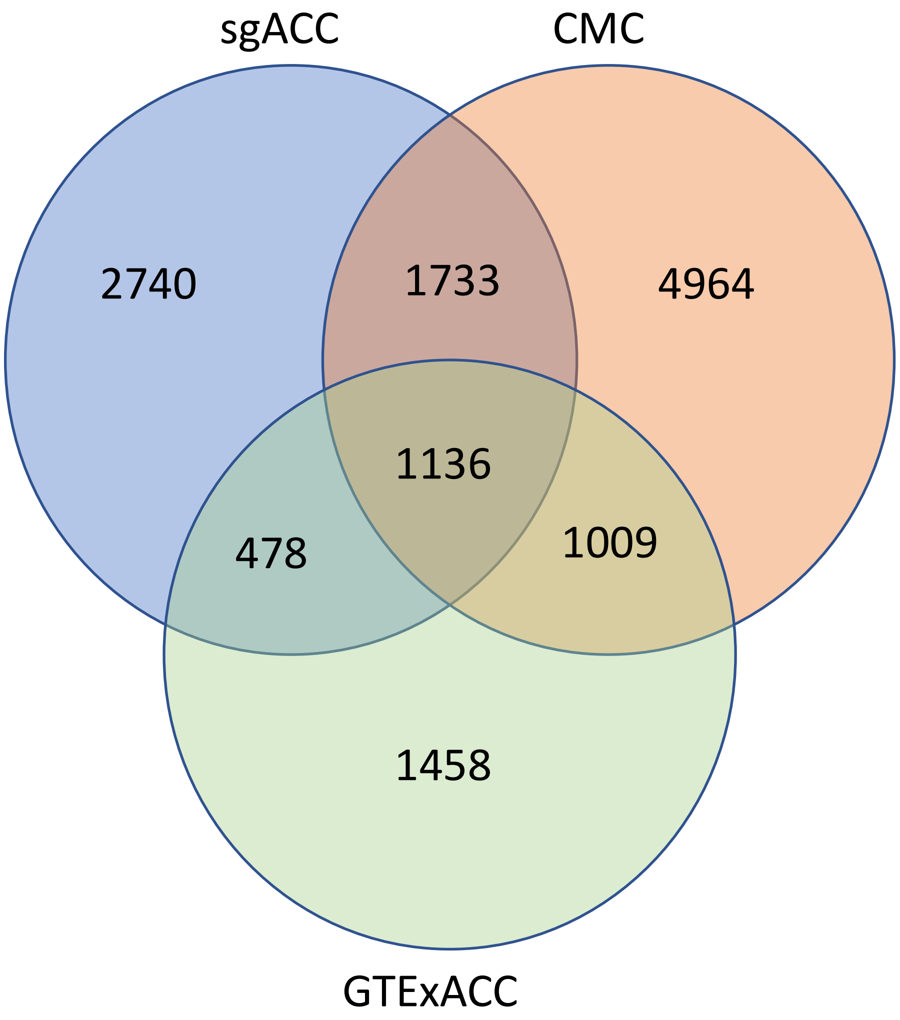

Supplementary Figure 6: Read count distribution of transcripts differentially expressed in any disorder (FDR<5%)

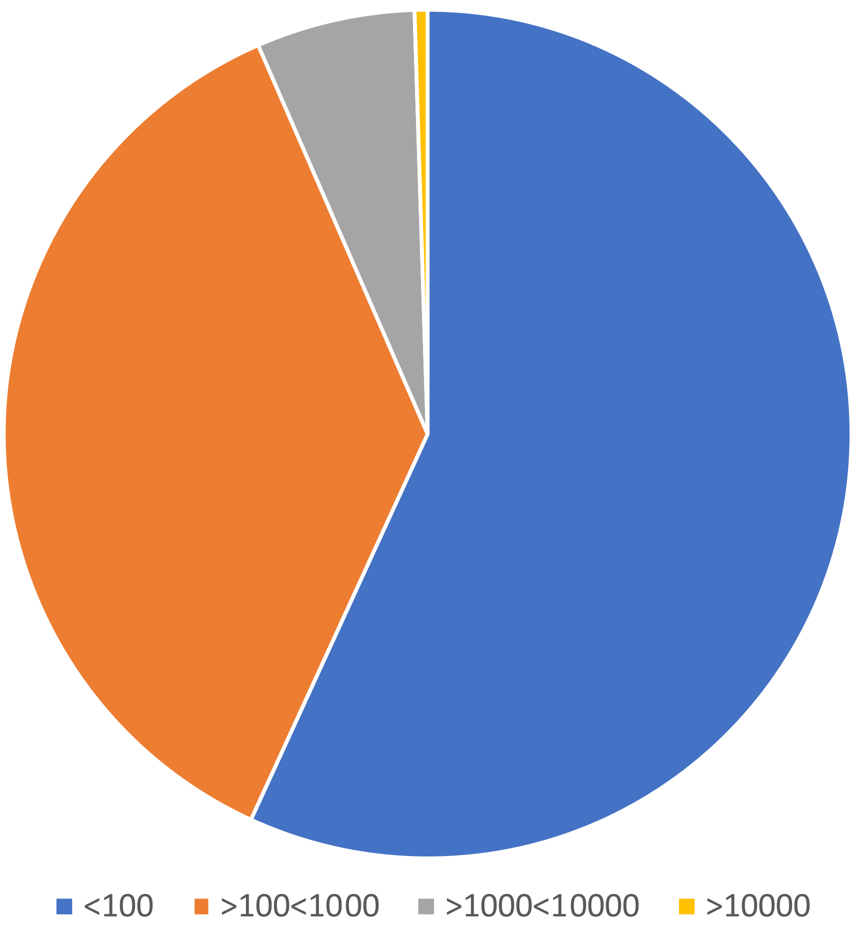

Supplementary Figure 7: Distribution of predicted alternative splicing events in subgenual anterior cingulate cortex (Caucasian sQTLs p<0.05)

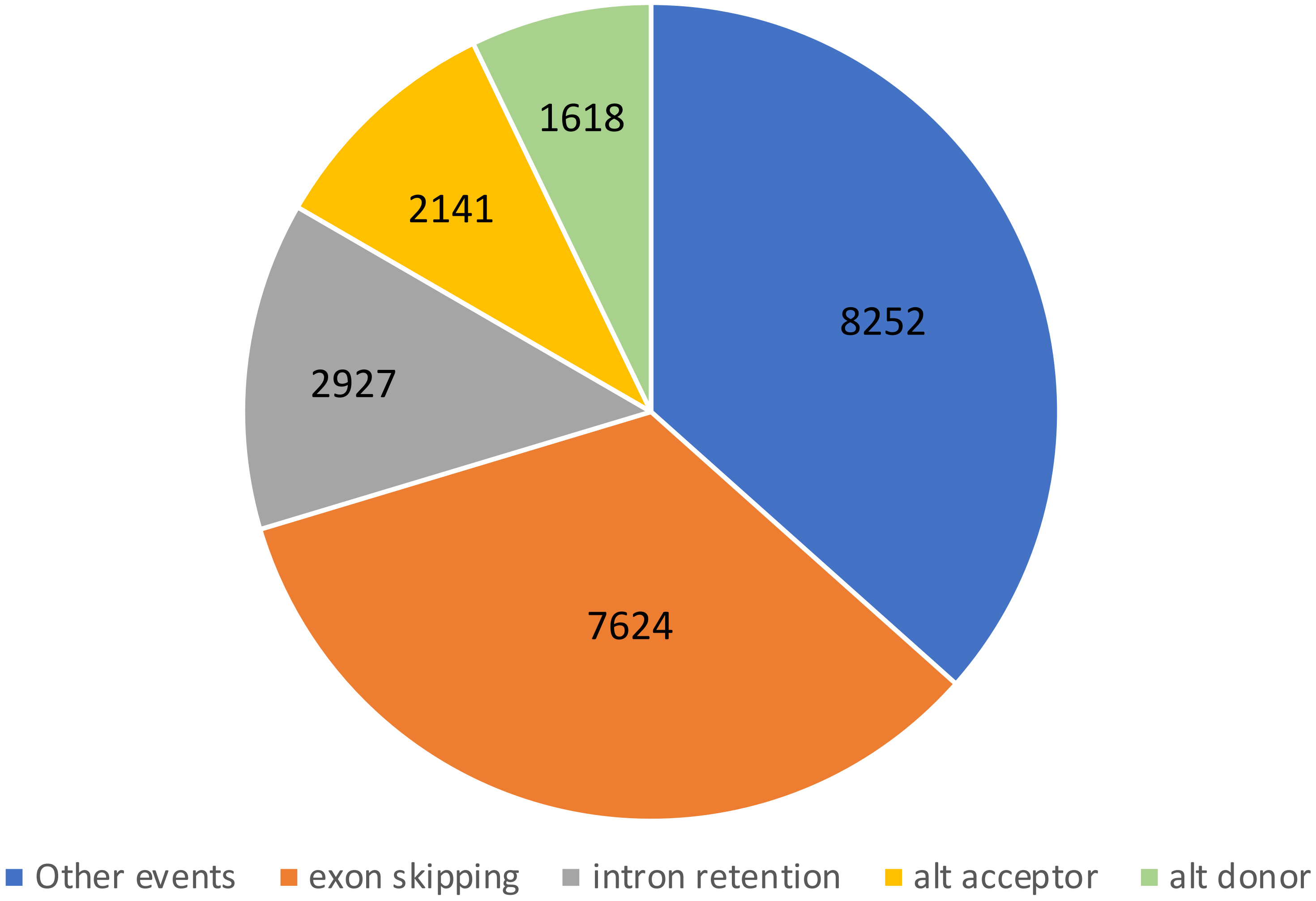

Supplementary Figure 8: Histogram of distances between sQTLs (p<0.05, Caucasian-only) and nearest splicing site

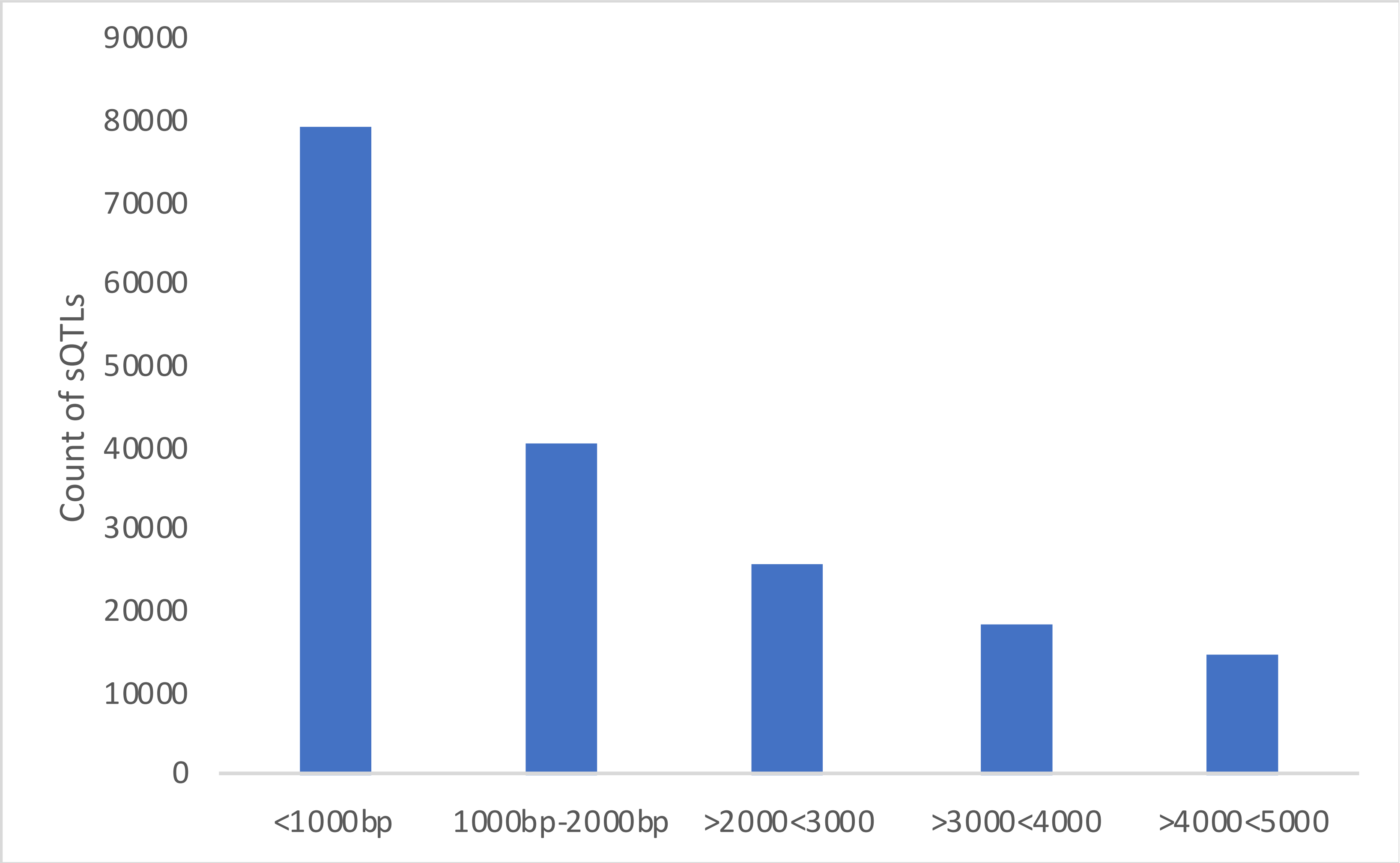

Supplementary Figure 9: Box plots of sQTL effect on relative transcript abundance. The two transcripts that change the most by a given variant (as identified by sQTLseekeR) are shown. If the transcript is differentially expressed vs controls, the diagnostic group is indicated above the boxplot for that transcript. The genotypes are shown on the x-axis and the relative transcript abundance is shown on the y-axis.

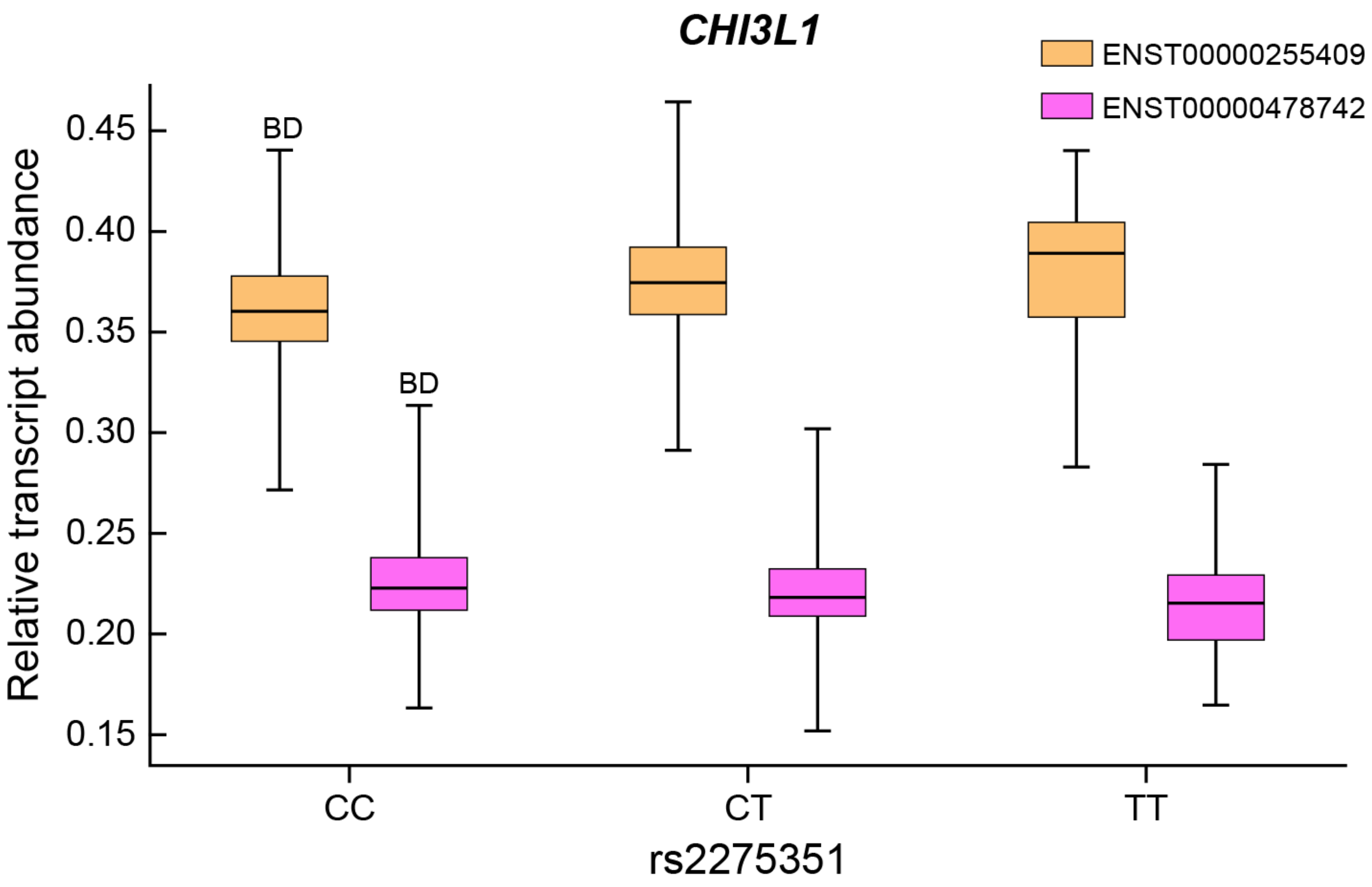

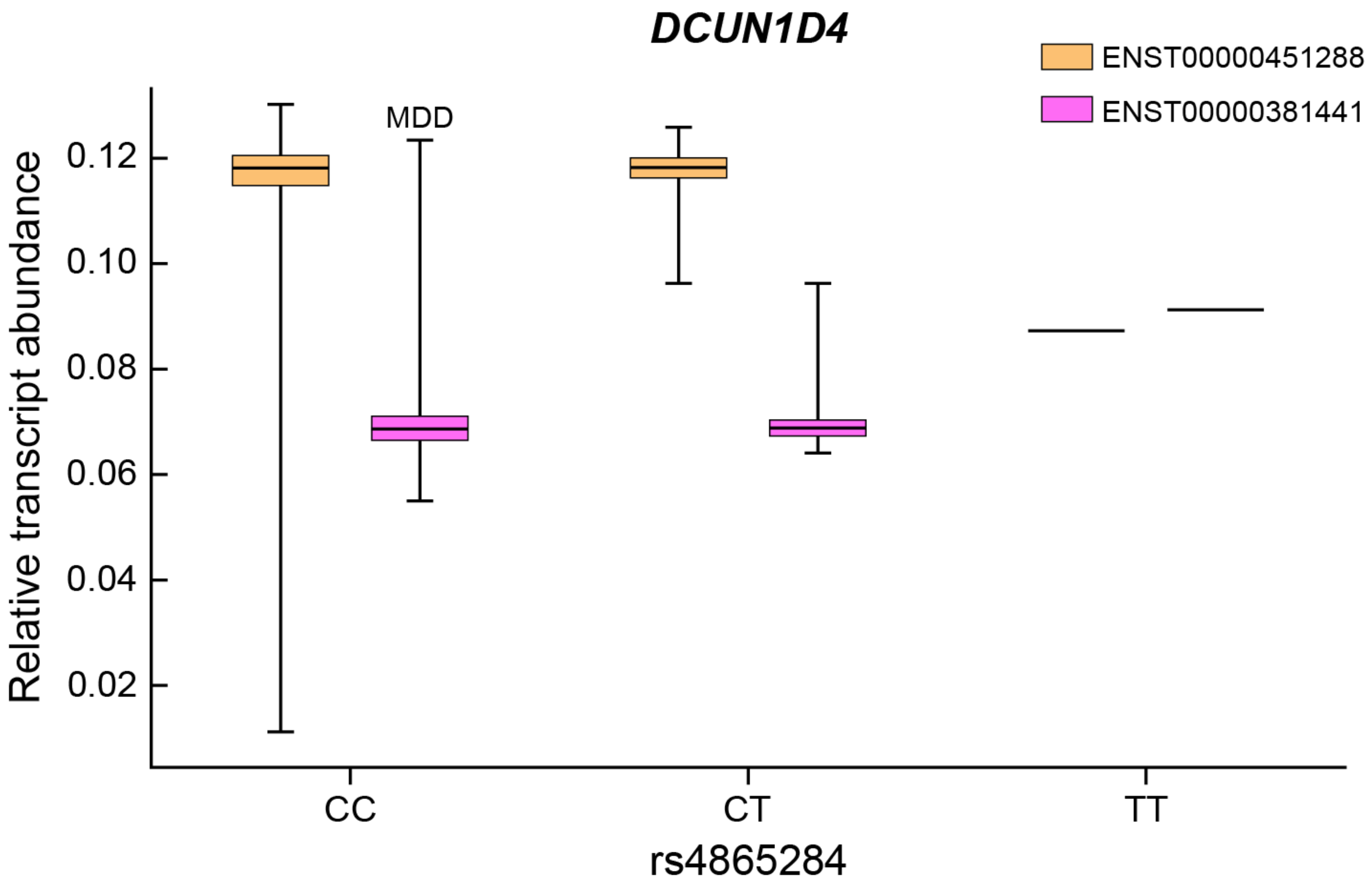

DCUN1D4
ENST00000451288
ENST00000381441
MDD
Relative transcript abundance
0.12
0.10
0.08
0.06
0.04
0.02
CC
CT
TT
rs4865284

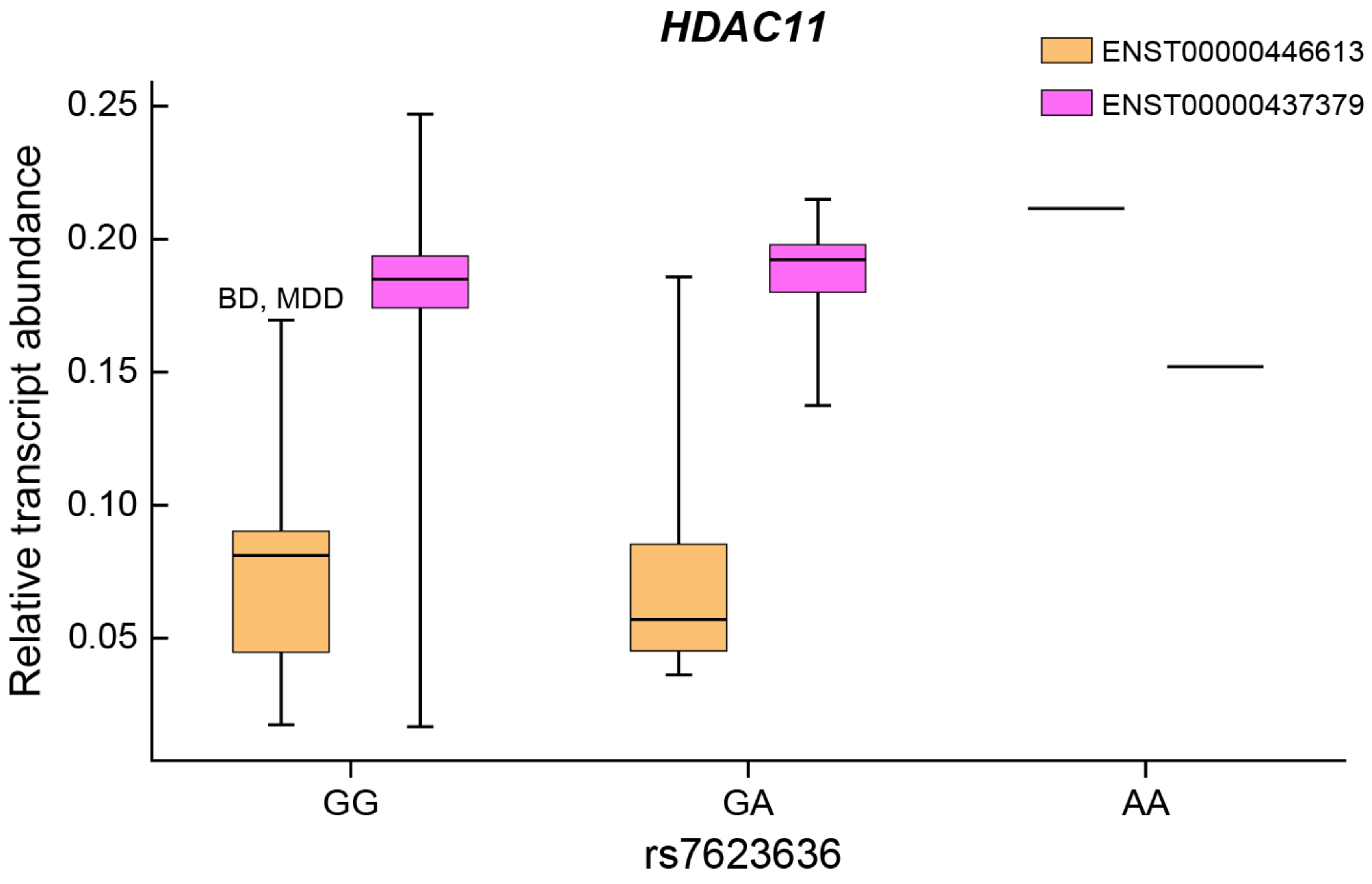
HDAC11
ENST00000446613
ENST00000437379
BD, MDD
Relative transcript abundance
0.25
0.20
0.15
0.10
0.05
GG
GA
AA
rs7623636

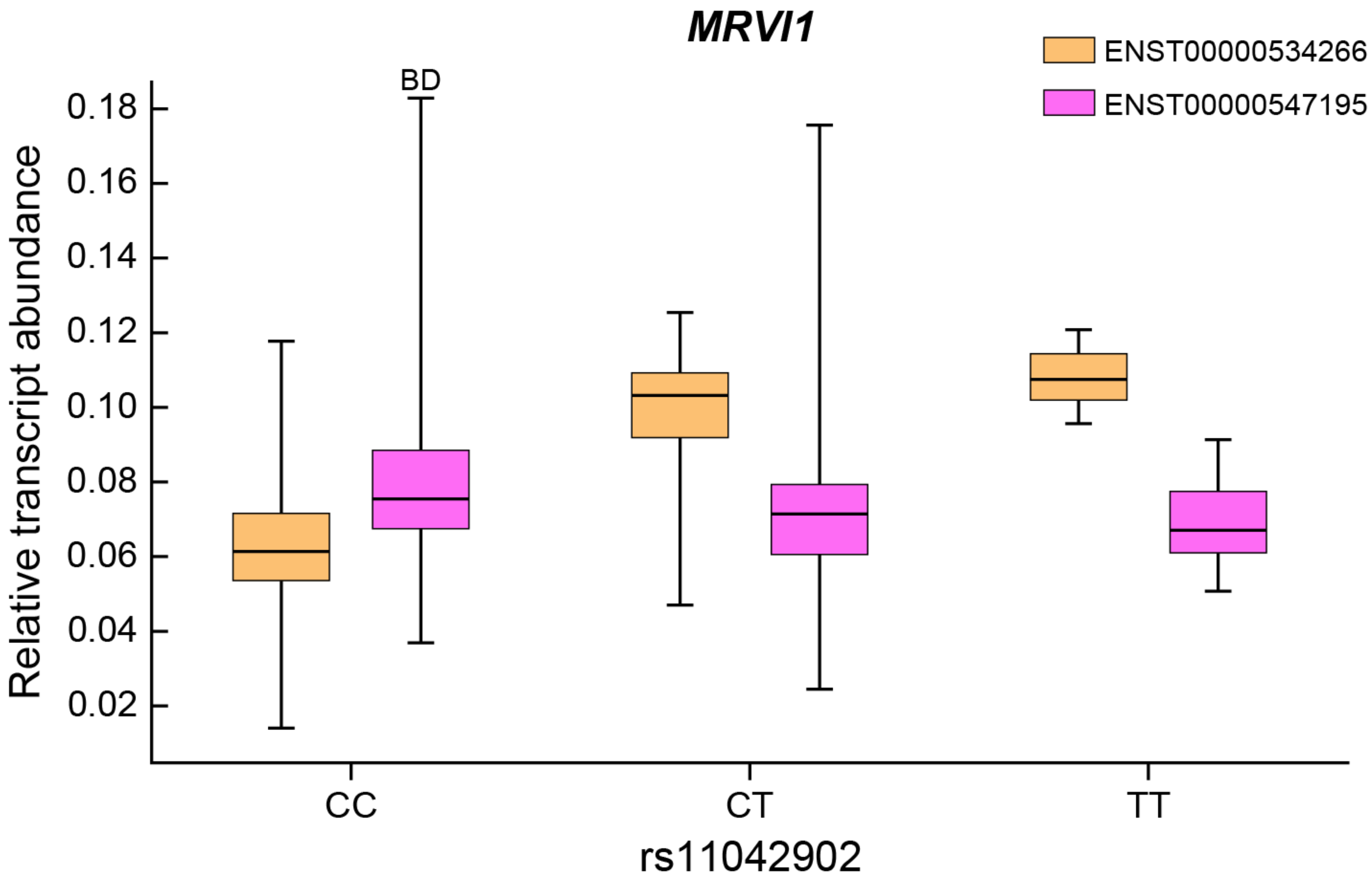
MRVI1
ENST00000534266
ENST00000547195
BD
Relative transcript abundance
0.18
0.16
0.14
0.12
0.10
0.08
0.06
0.04
0.02
CC
CT
TT
rs11042902

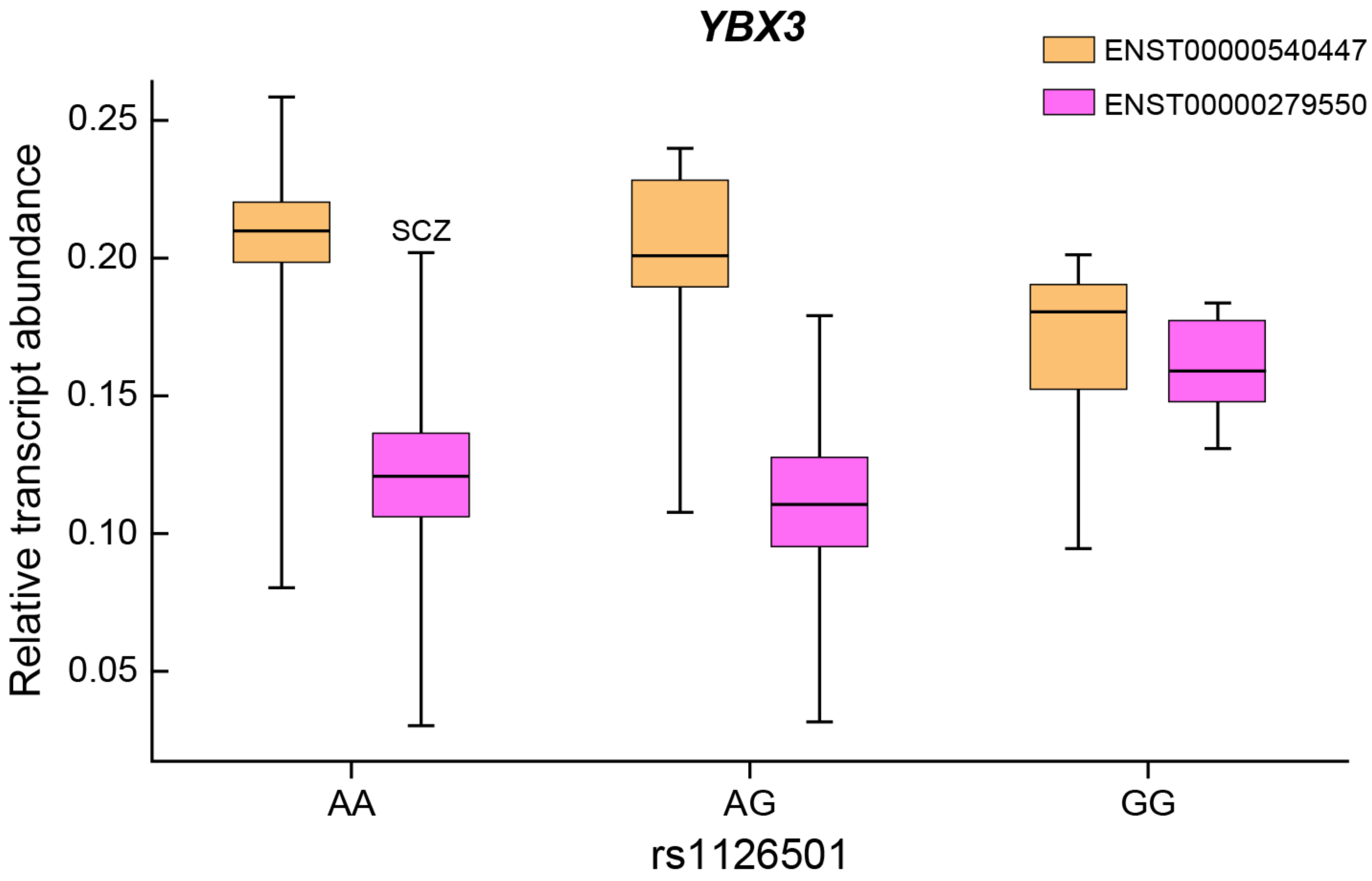
YBX3
ENST00000540447
ENST00000279550
SCZ
Relative transcript abundance
0.25
0.20
0.15
0.10
0.05
AA
AG
GG
rs1126501

Supplementary Figure 10: Correlation matrix of log2(FPKM+2) values, before and after removal of outlier samples

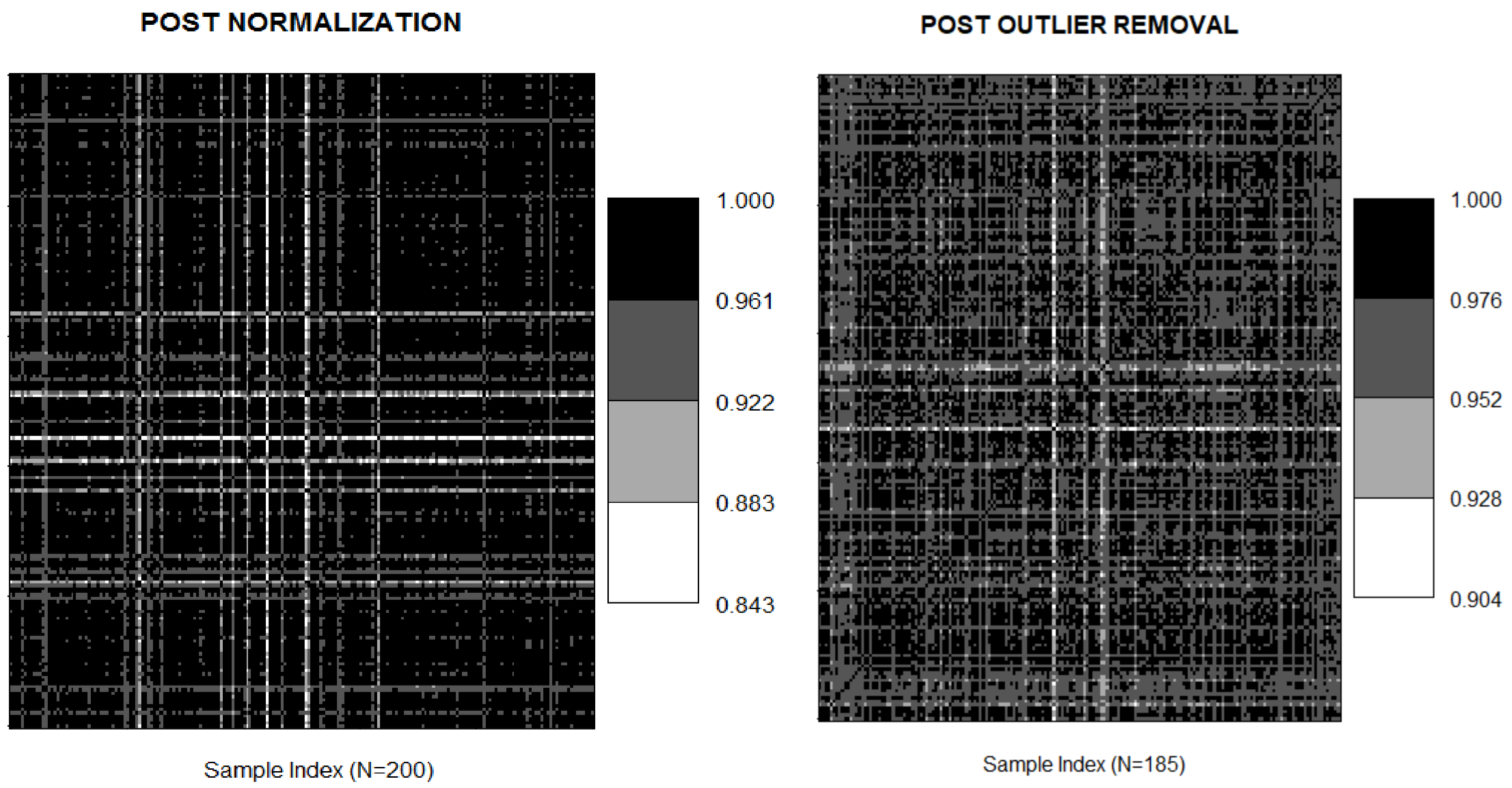

Supplementary Figure 11: Noise modeling of gene-level expression. Full distribution of counts is shown left. Insert shows zoomed region of plot shown on right.

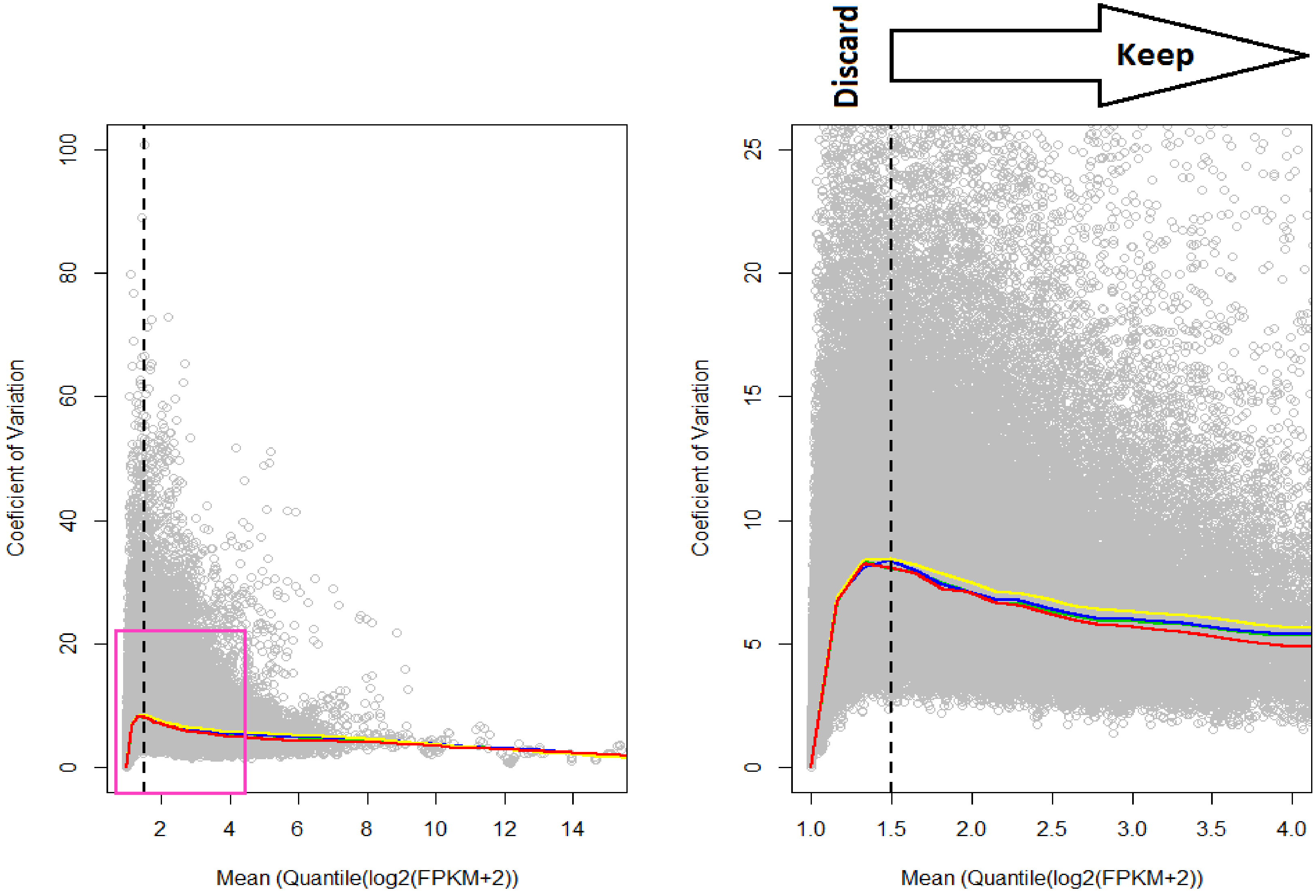
